# Solution Structure of the Novel CH-domain zinc finger from the puberty regulator Makorin-3

**DOI:** 10.64898/2025.12.01.691648

**Authors:** Antonio J. Rua, Andrei T. Alexandrescu

**Author notes:** Address correspondence to: Andrei Alexandrescu, Department of Molecular and Cell Biology, University of Connecticut, 91 N. Eagleville Rd., Storrs, CT 06269-3125, USA.

## Abstract

The makorin (MKRN) family of E3 ubiquitin ligases (MKRN1–4) regulates diverse biological processes, including reproduction, neurogenesis, and immune function. The first identified member, MKRN3, is an inhibitor of sexual development that is the site of inherited mutations linked to central precocious puberty (CPP). All makorin proteins share a distinctive cysteine/histidine-rich (CH) domain that has not been previously characterized experimentally. In MKRN3, the CH domain lies between the second C3H zinc finger and the RING domain. The C3H(2)–CH–RING segment appears particularly sensitive to CPP mutations suggesting it may constitute a structure-function unit. Using CD and NMR spectroscopy, we show that the CH domain folds upon coordination of a single Zn^2+^ ion with picomolar affinity. Spectroscopic and NMR pH-titration analyses identify a CCHC-type metal-binding site, typical of zinc fingers with protein-interaction functions. The NMR structure reveals the CH-domain adopts a canonical ββα zinc finger fold, despite atypical ligand spacing and the absence of conserved hydrophobic residues that usually stabilize this type of motif. Thermal denaturation monitored by multiple spectroscopic probes indicates sequential unfolding, with side-chain packing disrupted near 33 °C but zinc-stabilized secondary structure persisting to ∼63 °C, consistent with a molten-globule intermediate at high temperature. The function of the CH-domain remains unknown, but it could play a role in allosterically transmitting information on the RNA-bound state of the preceding C3H(2) domain to the subsequent RING domain. Based on a similar metal ligand spacing to a zinc finger from the protein FAAP20 and AlphaFold modeling, the CH-domain may have a ubiquitin-binding function, but this will need to be verified experimentally as AlphaFold also confidently predicts complexes with unrelated random proteins.

## 1. Introduction

The makorins (MKRNs) are a family of four E3 ubiquitin-protein ligases (MKRN1-4) involved in reproduction and other processes, including viral defense (MKRN1), neurogenesis (MKRN2), inflammation (MKRN2), and the immune response (MKRN4) [1–3]. The family likely evolved through duplication and retroposition of an ancient gonad-specific maternal-effect gene [4]. The first member discovered MKRN3 (also known as ZNF127), is a single-exon ubiquitin E3-ligase that is maternally-imprinted in several tissues through gene methylation so that only the paternal allele is expressed [5]. MKRN3 is a master regulator of sexual development, acting as an inhibitor of puberty [1,3,6]. Through mechanisms not fully understood, MKRN3 suppresses the release of gonadotropin-releasing hormone (GnRH), putting a “brake” on the hypothalamic-pituitary-gonadal (HPG) axis that activates puberty [1,7–10]. MKRN3 is highly expressed in the hypothalamic arcuate nucleus that makes several critical regulators of GnRH secretion during infancy, but MKRN3 levels decline at the onset of puberty in both mice and primate models [11,12]. Consistently, MKRN3 knockout mice show elevated GnRH and accelerated puberty [13]. MKRN3 was formerly listed as an understudied rare disease protein under NIH/NCATS program RFA-TR-22-030.

Much of the current information on MKRN3 function comes from genome-wide association (GWAS) studies of mutations linked to central precocious puberty (CPP) [6,14,15], a rare inherited disease affecting 1/10,000 children with a 10-fold higher frequency in girls [16,17]. Untreated CPP leads to accelerated puberty, reduced adult height, societal problems, psychological distress [18,19], and long-term risks including obesity, heart disease, stroke, type 2 diabetes, and estrogen-dependent cancers [16,19]. CPP can be largely managed through intramuscular injections of GnRH or implants that secrete the hormone [17,19,20] but these therapies can have unwanted side-effects such as weight gain [21], so multiple treatment options are desirable. While most occurances are idiopathic, 10-30% of familial CPP cases are associated with mutations in MKRN3 [16,20], the genetic locus with the highest percentage of CPP-linked mutations [6,22]. The MKRN3 mutations are clustered in a contiguous region between the second C3H zinc finger (ZNF) and the RING domain [15], we shall refer to as C3H(2)-CH-RING, demarcating this segment with putative roles in RNA binding and protein ubiquitination as critical to function.

There are two main ways MKRN3 is thought to suppress the onset of puberty – through regulation of mRNA transcription and through the ubiquitin degradation pathway [6]. In the arcuate nucleus of the hypothalamus, MKRN3 inhibits the *KISS1* and *TAC3* promoters for the genes encoding the secretagogues kisspeptin and neurokinin B, that trigger GnRH release [23]. There is also evidence that MKRN3 may affect GnRH levels [3]. It is unlikely MKRN3 binds promoters directly since the protein is not found in the nucleus [3] and its ZNF domains are not typical of DNA-binding motifs. Rather, MKRN3 could affect transcription through protein-protein interactions or through its ubiquitination of transcriptional regulators. While its C3H ZNFs suggest RNA-binding potential, RNA-binding has not yet been demonstrated for MKRN3 [3], unlike its homologs MKRN1 [24] and MKRN2 [25]. As far as its ubiquitination activity, MKRN3 works in conjunction with the E2 conjugating enzyme UBCH5A to ubiquitinate targets such as MBD3 (methyl-CpG-DNA binding protein 3) [26] and poly-A binding protein PABC1 [27,28].

MBD3 epigenetically silences GnRH, which triggers the HPG axis to start puberty [26,29].

PABC1 regulates the stability and translation of GnRH mRNA. Thus, MKRN3 acts as a brake on GnRH release and puberty in childhood. The dysregulation of MKRN3 through self-ubiquitination due to familial mutations could be one of the factors that lead to CPP [26,29].

Structural studies would aid the understanding of function but to date none of the four MKRN proteins have been structurally characterized. The sequences of the four MKRN proteins have a unique array of ZNF domains consisting of two or three C3H domains, a cysteine/histidine-rich CH domain unique to the makorin family, a RING domain, and final C3H domain [6,24,30,31]. Except for these domains the proteins are predicted to be intrinsically disordered in the absence of binding partners. The domain organization for MKRN3 is shown in Fig. 1A and a sequence alignment of the CH domains from the four makorins is given in Fig. 1B. The C3H domains are likely RNA-binding modules [2,5]. RING ZNFs are often the ‘active sites’ of E3 ubiquitin ligases, mediating the transfer of ubiquitin from an E2 ligase to the substrate destined for ubiquitination [2,32]. The CH-domain with three cysteines and three histidines [2,5,33] that is emblematic of MKRNs could be a ZNF since it has several potential metal ligands, although metal-binding for this domain has not yet been demonstrated [33].

**Figure 1.**
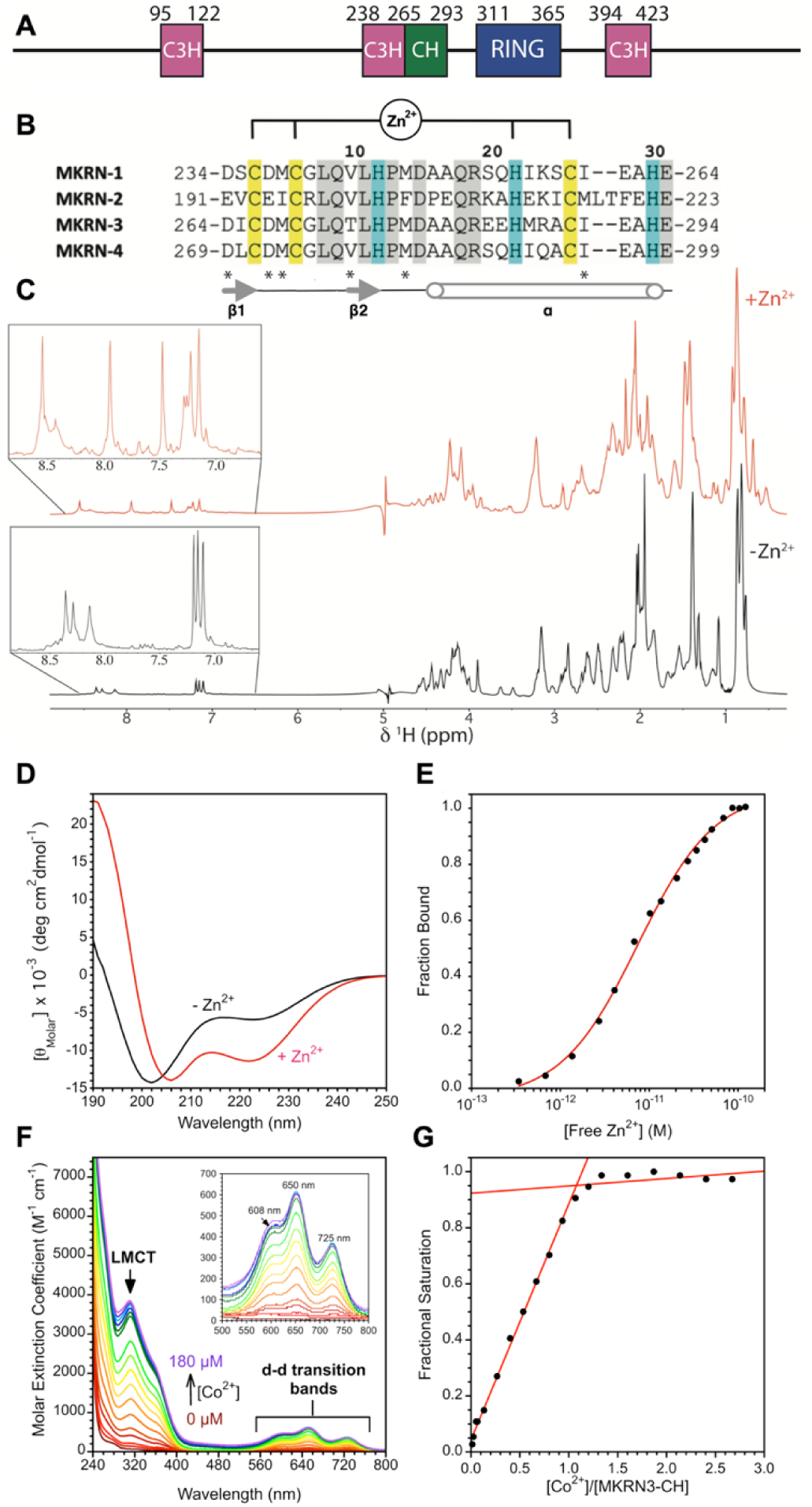
S**e**quence **and metal binding properties of the MKRN3-CH domain.** (**A**) Domain organization of full-length MKN3. (**B**) Sequence alignment of the CH domains from the four makorins. Cys (yellow), His (cyan) and conserved residues (gray) are highlighted. Positions where residues with similar properties occur are indicated by the symbol *. The four residues that ligate Zn^2+^ in the CH-domain are indicated at the top of the figure (see also Fig. 2). (**C**) 1D ^1^H NMR spectra of 1mM MKRN3-CH in D_2_O showing folding of the peptide in the presence of equimolar ZnCl_2_. (**D**) Zn^2+^-binding of MKRN-CH in the presence of the competitive chelator EGTA monitored by CD ellipticity at 222 nm. Analysis of the binding data as previously described [42,45] was used to determine a K_d_ of 7.5 x 10^-12^ M for Zn^2+^ binding by MKRN3-CH. (E) CoCl2 titration of a 0.75 µM sample of MKRN3-CH. Spectra are shown on a color ramp from red to violet going from low to high Co2+ concentrations, respectively. The inset is an expansion of the d-d region of the spectra (500 to 800 nm). Kd analysis of the data is given in Fig. S1. (F) A plot of fractional saturation of MKRN3-CH by Co2+ monitored by A650 indicates a 1:1 binding stoichiometry.

Given that the CH domain has an unknown function and a novel sequence pattern for a putative ZNF, it is important to determine if it folds upon binding zinc. The UniProt database [34] (accession code: Q13064) does not designate the MKRN3-CH region as a ZNF, and the ZF server (https://zf.princeton.edu/logoMain.php) [35] fails to detect a ZNF in the CH-domain sequence. As was the case with our recently reported ZC4H2 example [36], AlphaFold3 [37] (AF3) predicts with high-confidence (pLDDT > 70) folded models for 80% of a dataset of 30 random amino acid sequences constrained to have the same 31-residue length and 3Cys+3His content as the MKRN3 CH-domain sequence. Moreover, 87% of the random sequences, which are unlikely to fold, are predicted to have four or more of their cysteines and histidines within bonding distance of the zinc atom. Thus, AF3 is unreliable for predicting whether the CH domain binds metals or is folded. Some authors have inferred the CH domain is a CCCH ZNF [10] whereas others have suggested a CCHC ligand set [14,38]. Yet its sequence is not a good match for either. CCCH ZNFs like the ones that occur in MKRNs typically have RNA-binding functions and a motif with the sequence spacing C-X_7-10_-C-X_4-5_-C-X_3_-H [39], although exceptions are known [40]. The closest match in the MKRN3-CH sequence (Fig. 1B) would be C-X_2_-C-X_19_-C-X_3_-H. There are two types of CCHC ZNFs, a C-X_2_-C-X_4_-H-X_4_-C motif associated with retroviruses and RNA binding [39,41], and a C-X_2-5_-C-X_12_-H-X_3-5_-C motif associated with protein-protein interactions [42,43]. The closest match in the MKRN3-CH sequence (Fig. 1B) is C-X_2_-C-X_15_-H-X_3_-C. Hence the CH domain of the MKRNs has a unique sequence pattern, supporting its classification as a novel motif. A previous study inferred an all α-helical structure for the CH-domain using homology modeling approaches but model coordinates were not reported [38]. To obtain a structural characterization we synthesized a 264-294 peptide fragment of MKRN3 that encompasses the CH domain according to the UniProt database (accession code Q13064, residues 266-293). We show using CD and NMR spectroscopy that the fragment adopts a folded structure upon binding zinc. NMR pH titration experiments on histidines and UV-Vis of the peptide complexed with the spectrophotometrically active metal Co^2+^ were used to identify the residues that ligate Zn^2+^. The *K*_d_ value for Zn^2+^-binding was determined using a CD assay. We show that thermal unfolding of the Zn^2+^-bound CH-domain gives non-coincident transitions by different spectroscopic probes, suggesting denaturation of the metal binding site precedes complete unfolding of the secondary structure. The present results establish the CH-domain of MKRN3 as a bona-fide ZNF, findings that should be generalizable to the homologous CH domain in the other members of the MKRN protein family.

## 2. Materials and methods

### 2.1. Peptide and reagents

The 31-amino acid MKRN3-CH peptide, corresponding to fragment 264-294 of the native MKRN3 sequence, was custom synthesized at 90.2% HPLC purity by Biomatik (Kitchener, Canada). The peptide had its termini blocked by N-acetylation and C-amidation to more closely resemble the corresponding segment with no free charged ends in the full-length protein. The molecular weight by mass spectrometry was within 0.1 Dalton of that expected from the sequence. Peptide concentrations were determined using the BCA assay [44]. D_2_O (99.9% ) was from Cambridge Isotopes (Tewksbury, MA). ZnCl_2_ (purity 97%), CoCl_2_ (purity 99%), and all other reagents were from Sigma (St. Louis, MO).

### 2.2 Circular dichroism experiments

Circular dichroism (CD) data were obtained on an Applied Photophysics Chirascan V100 Spectrometer (Surrey, UK) employing a 1 mm path-length cuvette. Spectra were recorded between 190 and 250 nm, with a 1 nm bandwidth, a 1 nm step size, and 5 s/point data averaging, resulting in total experiment time of ∼5 min per spectrum. All CD experiments except for the temperature unfolding experiments were done with peptide samples in 10 mM NaPO_4_ buffer, pH 7.0, containing 0.2 mM of TCEP (tris(2-carboxyethyl)phosphine) to maintain cysteines in their reduced state. A 75 µM peptide concentration was used to compare spectra of the CH-domain with and without 90 µM ZnCl_2_. The *K*_d_ value for Zn^2+^-binding was obtained according to a published assay [42,45] utilizing the competitive chelator EGTA (17.71 mM), with 124 µM CH-domain peptide and a 0-350 µM range of ZnCl_2_ concentrations. Free zinc concentrations and the *K*_d_ value were calculated as previously described [42,45,46]. Temperature unfolding of the Zn^2+^-and Co^2+^-bound CH-domain by CD was investigated using 10 mM NaPO_4_ buffer, pH 6.0, containing 0.2 mM of TCEP with 100 µM peptide in the presence of either 120 µM ZnCl_2_ or 120 μM CoCl_2_. The temperature was ramped from 7 to 96 in 10-degree steps before being brought back down to 7 in 10-degree steps to check reversibility. Spectra were obtained with the CD parameters above, but with 0.75 s/point data averaging, to shorten the scan time to 72 seconds per temperature point. After the temperature ramp, a spectrum at 20 °C verified the changes in CD with temperature were reversible.

### 2.3 UV-Vis spectrophotometry of cobalt complexes

Binding of spectrophotometrically active Co^2+^ to the CH-domain was studied on an Ultrospec 8000 double-beam UV-Vis spectrophotometer (Thermo Fisher) with 1 cm pathway cuvettes. The MKRN3-CH peptide was 75 μM in 10 mM Tris buffer, pH 7.0, with 0.5 mM of TCEP. The CoCl_2_ was varied from 0 to 200 μM to obtain binding data. To determine the *K*_d_ value, we monitored the increase in the d-d band absorption at 650 nm (which is characteristic of tetrahedrally coordinated Co^2+^) as a function of CoCl_2_ concentration. The binding data were fit to the direct-metal titration equation:

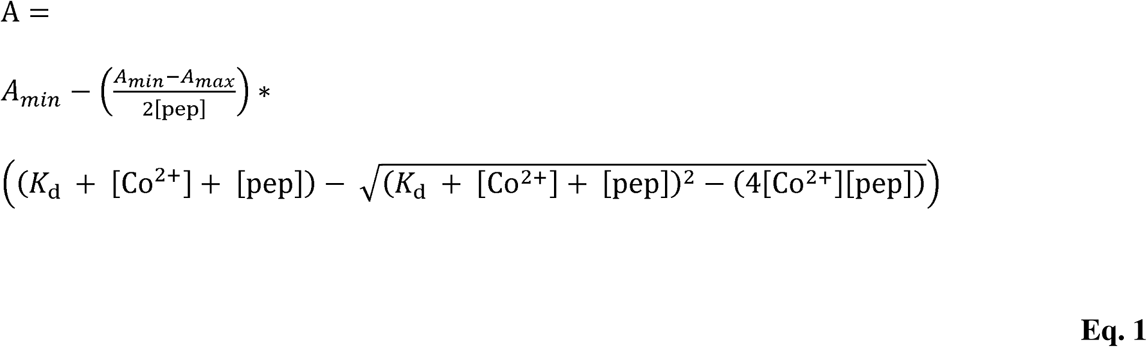

In the equation A_min_ and A_max_ are the limiting absorption values without and with saturating Co^2+^, and [pep] and [Co^2+^] are the respective total concentrations of peptide and metal [47,48].

To investigate the temperature-dependence of the d-d band from the cobalt complex, we collected 400-800 nm data on the Photophysics Chirascan V100 Spectrometer, operating in UV-Vis mode. The spectrometer was used because of its ability to sample data over a wide temperature range. Spectra were collected using a 1 cm cuvette, with 0.5 s/point data averaging, over a temperature range from 11 to 94 . Three spectra were averaged at each temperature for improved signal-to-noise. The sample had a 200 µM concentration of the CH-domain with 230 µM CoCl_2_, in 10 mM Tris, 0.5 mM TCEP, pH 7.

### 2.4 NMR spectroscopy

Unless stated otherwise, NMR experiments were done on a Bruker Avance 800 MHz instrument with a cryogenic probe at a sample temperature of 7 °C to avoid loss of amide protons due to solvent exchange. Samples had 1.9 mM MKRN3-CH in 270 μL of 90% H_2_O/10% D_2_O with a slight excess of 2.3 mM ZnCl_2_ at pH 6.0. The sample had no other added buffers or salts. 2D NOESY (200 ms mixing time), 2D TOCSY (70 ms mixing time), and a natural abundance ^15^N-SOFAST-HMQC spectra were collected for NMR assignments and structure determination. The latter ^15^N-SOFAST-HMQC was obtained on a Varian 600 MHz with a cryoprobe, at a higher temperature of 25 °C for improved sensitivity. A DOSY spectrum of this sample was also recorded on the Varian 600 MHz instrument at 25 °C, to avoid problems with thermal convection at other temperatures [49]. The sample was lyophilized and redissolved in D_2_O, to collect an 800 MHz natural abundance ^1^H-^13^C-HSQC spectrum at 7 °C. ^1^H and ^13^C were directly referenced to internal DSS (2,2-dimethyl-2-silapentane-5-sulfonate). ^15^N was indirectly referenced as described in the literature [50]. The extent of resonance assignments were: backbone ^1^H and ^15^N (100%), all ^1^H (83.71%), all ^15^N (74.42%), and all ^13^C excluding C’ (78.30%). NMR assignments have been deposited to the BMRB database under accession number 53190.

NMR temperature titrations were done on 1.9 mM MKRN3-CH containing 2.3 mM ZnCl_2_ in D_2_O at pH 6, on a 600 MHz Varian INOVA instrument equipped with a room temperature probe. NMR spectra were obtained both while increasing and decreasing the temperature to verify reversibility. The intensity of the Leu11 Hδa resonance was calibrated to the invariant intensity of the NMR signal from the DSS chemical shift reference standard, to obtain the folded fraction of the CH-domain at each temperature.

NMR pH titrations of a 0.8 mM zinc-bound MKRN3-CH sample were done in D_2_O at a temperature of 7 °C on a Varian 600 MHz instrument. The sample pH was monitored with a Mettler (Columbus, OH) MA235 pH meter equipped with a glass InLab Micro pH electrode at 25 °C, both before and after NMR experiments. The pH was calculated as the average of the two measurements. The p*K*_a_ values for titrating histidines were calculated from fits of their chemical shifts as a function of pH to a Henderson-Hasselbalch equation [51].

Finally, a 1.7 mM MKRN3-CH sample with 2.0 mM ZnCl_2_ in 90% H_2_O/10%D_2_O was used to collect a Bruker 600 MHz 70-ms mixing time 2D TOCSY experiment at a temperature of 7 °C and a sample pH of 10.8. Since H/D isotope exchange is too fast to measure in D_2_O for the zinc-bound MKRN3-CH peptide at an acidic pH of 6.0, we used survival of NMR signals in the 2D TOCSY at a high pH of 10.8, as an alternative to identify amide protons protected from solvent exchange for use in H-bond restraints for NMR structure calculations.

### 2.5 Calculation of NMR solution structures

Distance restraints for MKRN3-CH were obtained from the 200 ms mixing time NOESY experiment described above and were grouped into three broad distance ranges based on NOE intensities: 1.8-3.0, 1.8-4.0, 1.8-5.0 Å. Pseudoatom corrections to NOE upper bounds for prochiral atoms were included as described [43,52]. Torsional restraints were calculated from assigned ^1^HN, ^1^Hα, ^15^N, ^13^Cα and ^13^Cβ chemical shifts using TALOS-N [53].

The amino acids that coordinate Zn^2+^ were identified from the NMR pH titration and Co^2+^ binding experiments. Distance restraints from the three cysteines C3, C6, and C26 to Zn^2+^ were set to 2.33-2.37 Å and 3.25-3.51 Å for Zn^2+^-Sγ and Zn^2+^-Cβ, respectively. For H22, a restraint of 1.80-2.20 Å was set between the Nδ histidine atom and Zn^2+^, based on preliminary structures that showed a preference for the Nδ over the Nε histidine atom to ligate zinc.

Additional restraints between the C3, C6, and C26 Sγ atoms and the H22 Nδ atom were included to enforce the tetrahedral geometry of the coordination site, as previously described [54]. Hydrogen bond restraints were included for amide protons in secondary structure inferred to be protected from solvent exchange by H-bonding based on presence of their NMR signals in the TOCSY experiment obtained in 90% H2O / 10% D2O at pH 10.8.

Initial structures were calculated with a distance/geometry simulated annealing protocol in XPLOR-NIH [55]. A structure with no dihedral violations >5° and no NOE violations >0.5 Å was selected from the X-PLOR structures to be used as the template for further structure and water refinement using ARIA 2.3.2 [56]. An initial set of 200 structures were calculated with ARIA 2.3.2 on the NMRbox platform [57] with the iteration protocols of the program left at their default values. The 20 structures with the lowest energies and no violations above the specified thresholds were chosen for analysis and PDB deposition under accession code 9P2Q.

## 3. Results

### 3.1 The CH domain is a ZNF that folds upon binding a single metal ion

Figure1A shows the domain organization of MKRN3. The CH domain, which is the focus of this study, is located between the second of three C3H domains that typically serve as RNA-binding modules, and the RING domain that mediates transfer of ubiquitin from an E2 ligase to the substrate [58]. The CH domain, which is conserved among the four makorin family proteins (Fig. 1B) is presumed to be a ZNF due to its high content of cystines and histidines [33].

Fig. 1C shows that a synthetic peptide corresponding to the MKRN3 CH domain becomes structured upon binding Zn^2+^ by NMR. Folding of the CH domain is supported by the appearance of NMR signals below 0.7 ppm, typical of methyl groups shifted upfield by ring currents from aromatic residues, as well by the increased dispersion of resonances from the histidines between 8.5 and 7.0 ppm (expanded in the insets). The somewhat limited NMR dispersion for the folded CH domain is probably due to three histidines being the only aromatic residues, and the absence of aromatic residues at usually conserved ZNF hydrophobic core positions [42,59,60].

CD further supports Zn²⁺-induced folding of the CH domain, with spectra displaying minima at 208 nm and 222 characteristic of folded α-helical secondary structure (Fig. 1D).

Changes in the CD ellipticity at 222 nm were used to determine a *K*_d_ value for Zn^2+^ of (7.5 ± 0.4) x 10^-12^ M at pH 7.0 and a temperature of 21 °C (Fig. 1E). To probe metal coordination further, we examined binding of the CH domain to the spectrophotometrically active metal Co^2+^ (Fig. 1F). The ligand-to-metal charge transfer (LMCT) extinction coefficient for Co^2+^ at about 320 nm gives a value of about 900-1200 M^-1^ cm^-1^ for each bound sulfhydryl group [61]. The value of 3850 M^-1^ cm^-1^ for the CH-domain at saturating Co^2+^ concentrations indicates all three cysteines in the CH domain participate in metal coordination. The cobalt d-d transition region of the spectrum exhibits three bands between 605 and 725 nm. The extinction coefficients > 300 M^-1^ cm^-1^ indicate a tetrahedral metal coordination geometry [61], while the presence of three absorption bands at 608, 650, and 725 nm are consistent with three cysteine and one histidine ligand forming the coordination site [42,43,62]. A plot of fractional saturation as a function of [Co^2+^]/[MKRN3-CH] equivalents indicated a one-to-one metal-binding stoichiometry (Fig. 1G). The binding data were used to determine a *K*_d_ value of (1.8 ± 0.6) x 10^-6^ M for Co^2+^ binding to the CH-domain at pH 7.0 and a temperature of 25 °C (Supporting information Figure S1).

Finally, DOSY spectra for the CH-Zn^2+^ complex (Fig. S2) gave a diffusion coefficient (D = 0.97 x 10^-10^ m^2^ s^-1^) and radius of hydration R_h_ = 10.9 Å (normalized against the diffusion standard DSS, D = 3.17 x 10^-10^ m^2^ s^-1^, R_h_ =3.34 Å), consistent with a monomeric oligomerization state for Zn^2+^-bound MKRN3-CH [49]. Thus, we conclude the CH-domain is a genuine monomeric ZNF that binds a single Zn^2+^ atom in a tetrahedral coordination site formed by the three cysteine ligands and one histidine.

### 3.2 The CH domain has a CCHC zinc coordination site

To further characterize the nature of the Zn^2+^-coordination site we performed an NMR pH titration on the Zn-bound MKRN3-CH domain in D_2_O (Fig. 2). Of the three histidines in the sequence the NMR signals of H12 and H30 shift with pH, whereas that of H22 does not, identifying this histidine as the Zn^2+^-ligand. The pH titration results are fully consistent with the cobalt binding experiments (Fig. 1F) indicating a metal coordination site comprised of all three cysteines in the sequence and one histidine; namely C3, C6, H22, C26 using the numbering scheme of the CH peptide (Fig. 1B). The CH domain is therefore a CCHC-type ZNF with a C-x_2_-C-x_15_-H-x_3_-C ligand spacing closest to the C-x_2-5_-C-x_12_-H-x_3-5_-C spacing of the previously described “CCHC-finger” ZNF motif, of which ZNF750 is an example [63]. Either of the alternative histidines in the CH-domain would give unprecedented and presumably highly unfavorable ligand spacing arrangements, namely C-x_2_-C-x_5_-H-x_13_-C for H12 as a ligand, and C-x_2_-C-x_19_-C-x_3_-H for H30 as a Zn^2+^ ligand.

**Figure 2.**
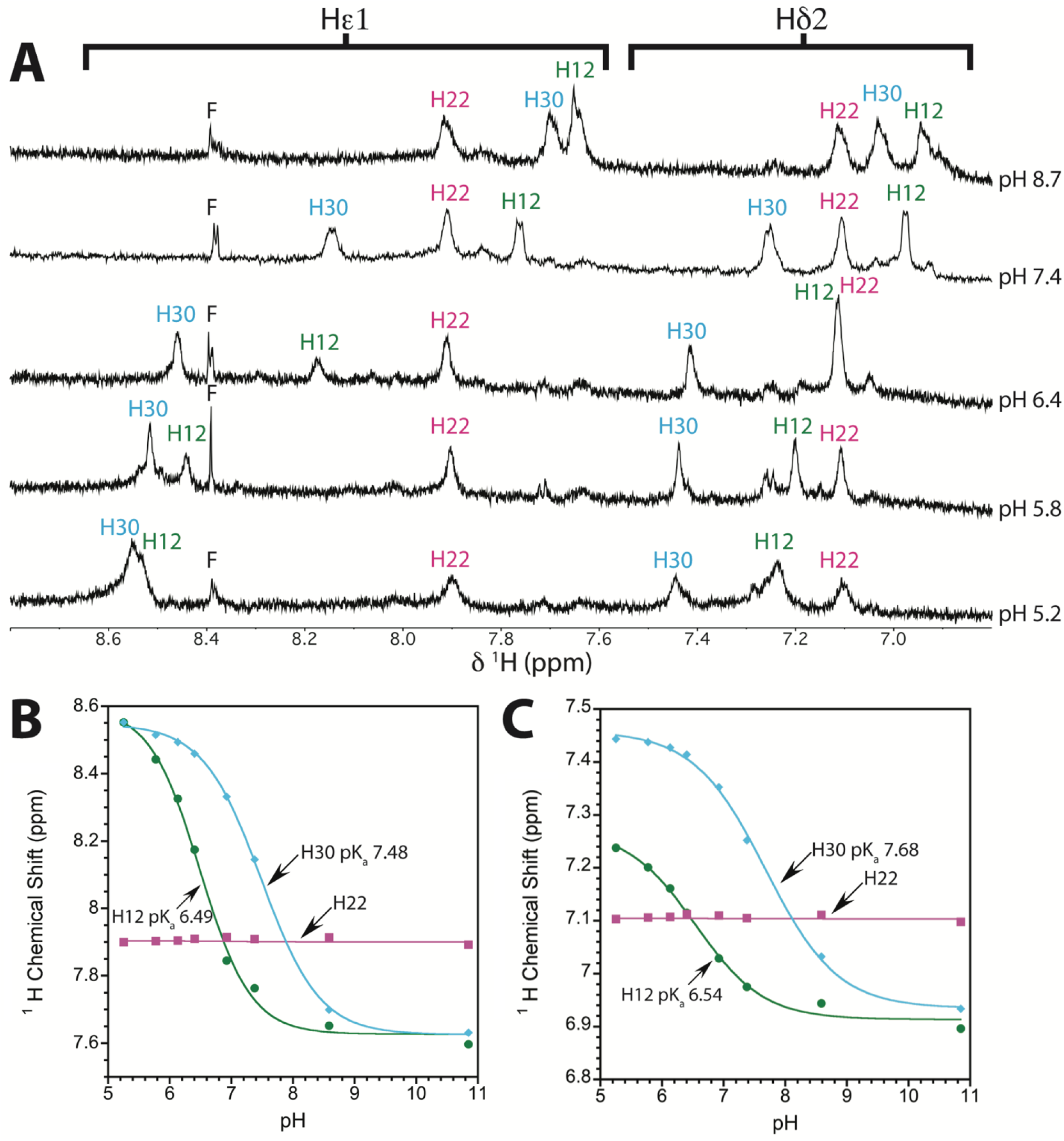
Identification of the zinc-ligating histidine in MKRN3-CH from an NMR pH titration. (A) Aromatic regions of selected 1D 1H NMR spectra of 1.9 mM Zn2+-bound MKRN3-CH in D2O as a function of pH*, showing NMR signals from histidine Hε1 and Hδ2 protons. Note that the three histidines are the only aromatic residues in the peptide. The resonance marked F is an impurity (formate). For the titrating His12 and 30, the data from the Hε1 (B) and Hδ2 (C) chemical shifts were fit to a modified Henderson-Hasselbach equation [51] to obtain the indicated microscopic pKa values. The Hε1 and Hδ2 resonances of His22 do not titrate, identifying this residue as the Zn2+-ligating histidine. All NMR data were obtained at a temperature of 7 oC.

The pH titration data gives p*K*_a_ values of 6.52 ± 0.04 and 7.58 ± 0.14 for H12 and H30, respectively. Thus, protonation of H30 should be more favorable than that of H12, presumably since H30 participates in favorable interactions in the CH-domain structure in its protonated charged state. By contrast, H22 resists protonation as this would interfere with Zn^2+^-binding, and hence this histidine does not titrate above pH 5.2, the most acidic pH we looked at (Fig. 2a).

### 3.3 The CH domain has a ββα structure

Having established the Zn^2+^-binding residues in the CH-domain we next turned to determining its solution structure by NMR. The necessary NMR assignments for structure determination were obtained as described in the Methods section. Figures 3A and B show ^13^Cα-^1^Hα and ^1^H-^15^N ‘fingerprint’ regions of natural abundance ^1^H-^13^C HSQC and ^1^H-^15^N SOFAST-HMQC spectra for the CH-domain, illustrating the completeness of the NMR assignments. NMR structures of the CH-domain were calculated from the restraints summarized in Table S1, that included NOE distance restraints, H-bonds identified from surviving of amide proton H-bond donors in high-pH NMR experiments, restraints to Zn^2+^ included from our previously described metal ligand identification (Fig, 2), and chemical-shift-derived torsion angle restraints calculated with the program Talos-N [53]. The set of 20 best NMR structures for the MKRN3-CH domain are shown in Fig. 3C. The CH-domain folds into the canonical ZNF ββα fold, despite the absence of any of the conserved aromatic and hydrophobic residues that typically stabilize this fold [42]. The CH domain gives backbone RMSDs of about 2.5 Å to both the CCHC ZNF from ZNF750 (PDB 8SXM [42]) and the CCHC ubiquitin-binding ZNF from FAAP20 (PDB 2MUQ, 2MUR [64]), which we found in a sequence motif search as the only ZNF that shares the ligand sequence spacing of the CH-domain (see Discussion). Thus, even though the CH-domain has sequence features that make it unrecognizable as a ZNF, its prototypical ββα fold is conserved.

**Figure 3.**
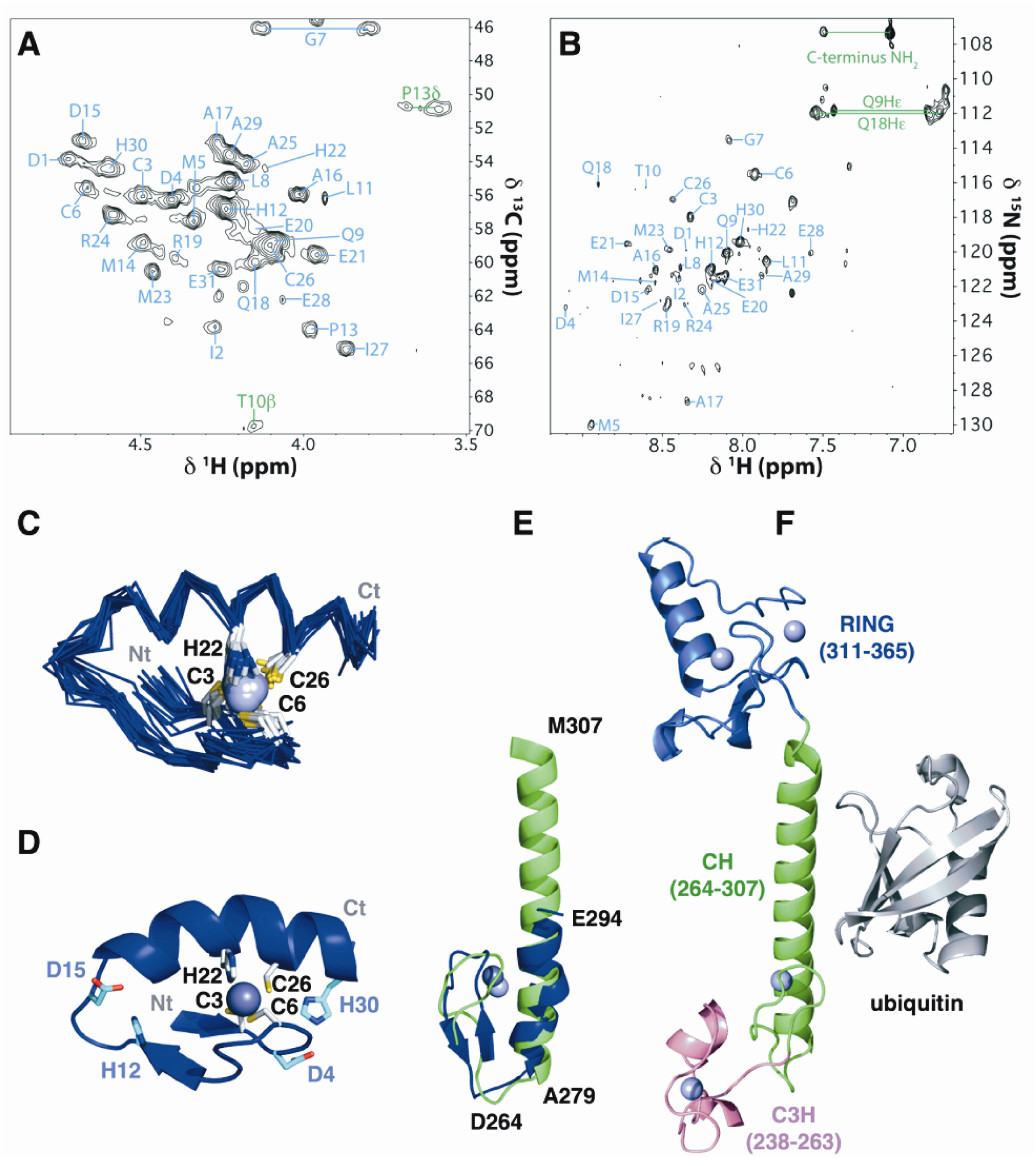
NMR assignments and structure of MKRN3-CH. Hα-Cα (A) and H-N (B) fingerprint regions of multiplicity-edited 1H-13C HSQC and 1H-15N SOFAST-HMQC spectra of the CH domain obtained at natural isotope abundance. (C) Bundle of 20 lowest-energy NMR structures for the CH domain. (D) Representative CH-domain NMR structure showing secondary structure, zinc-coordinating residues (white) and two ion-pair interactions (light blue). (E) Comparison of the CH-domain NMR structure (blue) and AF3 prediction (green). Note that the helix in the AF3 prediction extends past the CH domain and is twice as long as in the NMR structure. (F) AF3 prediction for full-length MKRN3 bound to ubiquitin, showing only the segment of MKRN3 encompassing the C3H(2)-CH-RING domains. The coloring scheme is the same as in Fig 1A. A space-filling diagram of interactions involving the CH-domain is shown in Fig. S6.

Fig. 3D shows a cartoon of a representative structure from the ensemble of CH-domain NMR structures, illustrating the position of the four zinc-coordinating residues C3, C6, H22, and C26. The two remaining non-coordinating histidines both appear to be involved in electrostatic ion-pair interactions in at least some of the NMR structures: H12 with D15, and H30 with D4.

Additional potential ion pair interactions include D1-R19, E20-R24, and E28-R24, with all of these occurring in a subset but not all of the NMR structures due to the limited precision of the sidechain conformers. A final potential stabilizing interaction is an H-bond in the interior of the structure between the side chain of Q18 and the backbone atoms of D1. These polar interactions are likely important in stabilizing the CH-domain structure since hydrophobic residues (I2, M5, L8, L11, M14, M23, I27) are located mostly on the surface surrounding strand β2 (L8, L11, M14), with only M5, L11, and partially M23 positioned towards the interior of the structure. The AF3 prediction for the CH domain similarly lacks buried hydrophobic residues.

Figure 3E compares the experimental NMR structure of the CH-domain (blue) with the corresponding segment from an AF3 prediction (green) of the full-length MKRN3 protein, modeled with 6 Zn^2+^ ions for the total number of zinc-binding sites in the protein. The experimental NMR and predicted AF3 structures are close with a backbone RMSD of 2.6 Å. A notable difference is that the α-helix in the AF3 prediction continues to twice the length (8 turns) of the experimental structure (4 turns). It is possible that the UniProt designated C-terminal domain boundary at H293 truncates the CH-domain too early. This explanation seems unlikely, however, as a truncation in the middle of an α-helix would typically disrupt the entire structure, not just a part. AF3 often predicts secondary structure for conditionally-folded intrinsically disordered regions [65], so the predicted α-helix extension may occur in the presence of a binding partner. Figure 3F shows the C3H(2)-CH-RING segment of MKRN3 from an AF3 prediction of the full-length protein in complex with ubiquitin (see Discussion).

### 3.4 The CH domain undergoes sequential thermal unfolding, with metal-stabilized secondary structure persisting after disruption of the tertiary structure

To assess the stability of the CH domain, given its unusual sequence and lack of a distinct hydrophobic core, we looked at the thermal stability of the metal-bound peptide using several spectroscopic probes. Fig. 4A shows selected points from a temperature titration of the Zn^2+^-bound CH-domain monitored by 1D ^1^H-NMR spectra. We were limited to temperatures between 7 and 49 °C for these experiments due to the temperature range of our NMR probe.

**Figure 4.**
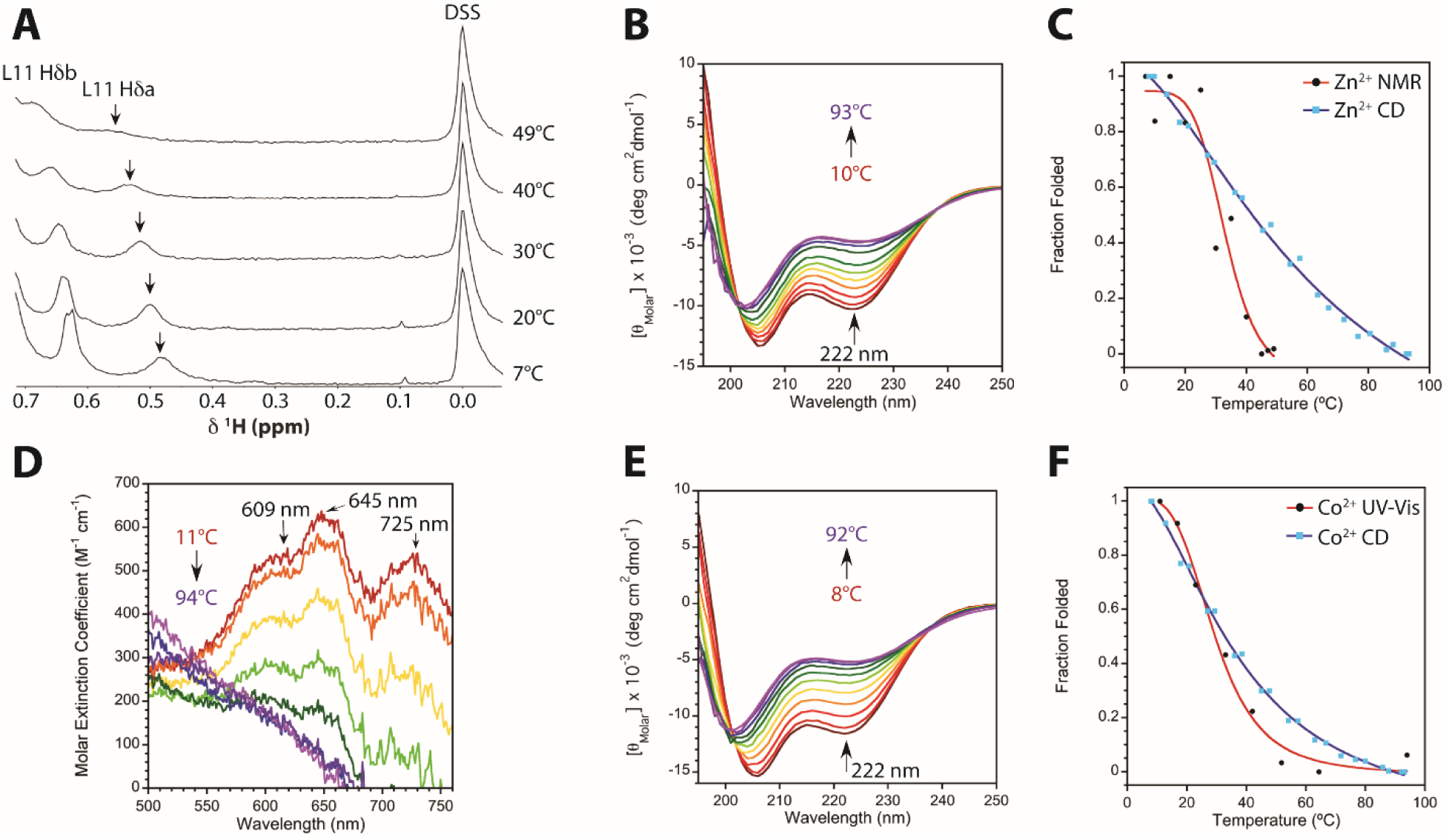
Stepwise thermal unfolding of MKRN3-CH. Temperature effects on (A) the high-field methyl region of 1D NMR spectra for Zn2+-bound MKRN3-CH. The complete spectra are shown in Fig. S4. (B) Temperature titration of Zn2+-bound MKRN3-CH monitored by far-UV CD. (C) Comparison of unfolding curves for the Zn2+-complexed CH domain calculated from the NMR data on tertiary structure in panel A (peak for L11 Hδa indicated by the arrow) and the CD data on secondary structure in panel B. Thermal unfolding of Co2+-bound MKRN3-CH by UV-Vis spectroscopy of the cobalt d-d transition region (D) and far-UV CD (E). The unfolding curves calculated from the UV-vis data reporting on metal coordination geometry and the CD data monitoring secondary structure are compared in (F).

The spectra show a decrease and broadening of NMR signals from high-field shifted methyl groups, suggesting exchange between folded and unfolded conformations in the slow to intermediate regime on the NMR timescale. For clarity, we show expansions of the spectra around the Hδa methyl signal from Leu11 that was used to quantify the fraction of folded protein. However, the broadening and loss of dispersed signals due to unfolding is seen throughout the 1D NMR spectra (Fig. S4).

Fig. 4B shows the thermal unfolding of the Zn^2+^-bound CH-domain by CD. The CD spectra show a gradual change with raised temperatures characterized by an increase towards a more positive ellipticity at 222 nm and a smaller increase in ellipticity together with a blue shift of the band at 208 nm towards 200 nm. We used the ellipticity at 222 nm to calculate the fraction of folded CH-domain. Small differences are seen between the CD-spectrum of the Zn^2+^- bound CH-domain at 93 °C (Fig. 4B) and the unfolded peptide in the absence of Zn^2+^ at 20 °C (Fig. 1D) indicating the CH-domain is not fully unfolded in the presence of zinc even at 93 °C. Namely, the ellipticity at 202 nm is slightly more positive at the higher temperature whereas the ellipticities at 222 nm are comparable. Temperature data on the apo CH-domain suggest these differences are temperature effects on the unfolded peptide (Fig. S5). A second factor in why the CD spectrum of the Zn^2+^-bound CH domain is not fully unfolded even at the highest temperatures, is that some residual structure persists even at 88 °C, as manifested by differences in the CD spectrum upon addition of ZnCl_2_ at high temperature(Fig. S5).

Figure 4C compares the thermal unfolding of the Zn^2+^-CH domain as assessed by NMR (intensity of the L11 methyl resonance) and CD (ellipticity at 222 nm). The two curves are markedly non-coincident, giving melting point (*T*_m_) midpoint values of 33 ± 4 °C by NMR and 63 ± 9 °C by CD. Non-coincident denaturation curves by multiple spectroscopic curves are a classic hallmark for sequential denaturation that proceeds through folding intermediates [66]. The NMR data on high-field methyl resonances is sensitive to the tertiary packing of sidechains in the core of the structure and shows a steep transition characteristic of a cooperative loss of structure. The far-UV CD data is sensitive to secondary structure and shows a shallow broad unfolding curve that is characteristic of ‘molten globule’ intermediates – conformational states that maintain loosely organized secondary structure but that lack the specific tertiary sidechain packing of native proteins [36,67,68].

We observe the same type of behavior in the thermal unfolding of the cobalt-bound CH-domain. For the Co^2+^-ligated CH domain we did not use NMR, as cobalt is a paramagnetic metal that broadens ^1^H-NMR signals. Rather, we looked at the temperature dependence of the d-d transition bands in the visible spectrum that are sensitive to metal coordination geometry (Fig. 4D) exhibiting extinctions coefficients > 300 M^-1^ cm^-1^, 50 to 250 M^-1^ cm^-1^, and < 30 M^-1^ cm^-1^ for tetrahedral, pentacoordinate, and octahedral coordination sites, respectively [47,61]. The decrease of the d-d absorption bands with temperature indicates a disruption of the tetrahedral coordination site for Co^2+^ but does not imply a loss of metal binding, since alternative coordination sites with a larger number of ligands have lowered extinction coefficients. Indeed, the data on the temperature dependence of the CD spectrum of the Co^2+^-bound CH-domain indicates secondary structure persists after denaturation of the tetrahedral coordination site for Co^2+^ (Fig. 4E). As we have shown throughout this work the secondary structure of the CH-domain is depended on metal binding. Taken together, the thermal denaturation data on the Co^2+^-bound CH domain is most consistent with a disruption of the tetrahedral metal coordination site to include additional ligands but the preservation of secondary structure underpinned by metal binding. Such a mechanism could occur if water penetrated the core of the domain at higher temperatures and bound to Co^2+^, resulting in pentacoordinate and octahedral coordination sites that result in decreased d-d band ellipticities, but that nevertheless provide a scaffold for the stabilization of the secondary structure of the domain. Like zinc, the transition curves for the cobalt-bound domain are non-coincident, although they are closer to each other, giving a *T*_m_ of 29 ± 2 °C for the UV-vis and 36 ± 1 °C for the CD data (Fig. 4F). The melting curve obtained for the metal coordination site from the d-d transition data shows greater cooperativity than from the unfolding of secondary structure by CD. The *T*_m_ for the melt of secondary structure by CD is much lower for the Co^2+^ (36 °C) than Zn^2+^ (63 °C) complex, consistent with a nearly six order of magnitude difference in the *K*_d_ values between the two metals (7.5 x 10^-12^ M for Zn^2+^ and 1.8 x 10^-6^ M for Co^2+^) (Fig. 1E, S1).

## 4. Discussion

Our work shows the MKRN CH-domain is a genuine ZNF with a CCHC Zn^2+^-ligand set typically associated with protein-protein interaction functions [36,42]. The CH domain is unusual in having an atypical 15-residue spacing between its second and third zinc ligands, and lacking the conserved position hydrophobic and aromatic residues that form a rudimentary hydrophobic core in ZNFs with a ββα-fold [42]. In fact, the CH-domain structure has only a few buried hydrophobic residues, which may account for its unusual unfolding properties. The Zn^2+^-bound CH domain shows noncoincident thermal unfolding transitions by different spectroscopic probes, a classical indicator for the presence of partially folded intermediates [66]. Loss of NMR signals typical of well-packed sidechains occurs at about 33 °C near physiological temperature, whereas secondary structure monitored by CD shows a shallow non-cooperative unfolding transition with an apparent midpoint of about 63 °C. Taken together, these results suggest that the CH-domain tertiary structure denatures near physiological temperature, but that the zinc-stabilized secondary structure persists in a molten globule-like conformation at temperatures some 30 °C higher. The same kind of behavior is seen for the Co^2+^-bound CH-domain, although the difference between the tertiary and secondary structure melting points is much smaller. The Co^2+^-bound form is not physiological, and we used it in these studies because the metal provides a sensitive probe of coordination sphere geometry through its d-d transition absorption bands [36,46,61]. With increasing temperature, the d-d transition bands indicate that the tetrahedral Co^2+^-coordination becomes denatured with a temperature midpoint of ∼29 °C, but that the secondary structure stabilized by alternative Co^2+^-coordination geometries with a larger number of ligating solvent molecules, has a higher temperature midpoint at about 36 °C.

The sequential thermal unfolding of the CH-domain through a molten globule intermediate that has metal-stabilized secondary structure but disordered side-chain packing, is consistent with our recent similar observations for the ZC4H2 zinc finger [36], and other theoretical and experimental results that support sequential folding of ZNF motifs from an initially partly-formed metal binding site [45,69,70]. That thermal unfolding is noncoincident by different spectroscopic probes cautions that multiple methods should be used to assess the viability and folding status of ZNFs. For example, the CH-domain would appear unfolded by NMR above physiological temperature but folded by far-UV CD. At the present time, we do not know if the disruption of sidechain packing in the CH-domain at physiologically temperature is biologically relevant. Our synthetic peptide construct could be omitting interactions from the full-length MK3 protein that destabilize the CH-domain at higher temperatures, although this seems unlikely. Our results indicate that the metal-induced secondary structure of the CH-domain can be uncoupled from its specifically packed sidechain tertiary structure.

Mutations in MKRN3 that lead to CPP are clustered to a three-domain segment C3H(2)-CH-RING, running from residues 238-365 in the full-length protein [15]. The clustering of the mutations suggests that this segment may be a functional unit, and this appears to be supported by AF3 modeling of intact MKRN3 (Fig. 3F, S6). We noted that the CH domain has few buried nonpolar residues, with most hydrophobic residues positioned on the surface near strand β2 in the CH-domain structure. The hydrophobic surface residues near strand β2 (residues L271-M277, yellow) are conserved and, according to the AF3 prediction, are positioned to interact with a hydrophobic patch (F243-R249, purple) from the α-helix between the first and second coordinating cysteines of the preceding RNA-binding domain C3H(2), as shown in Fig. S6A. Moreover, the AF3 prediction for full-length MKRN3 has the C-terminal α-helix from the CH-domain (A279-D308) running close to the start of the subsequent RING domain (V310-R365).

The significance of the C3H(2)-CH-RING domain organization is that the central CH domain could act as an allosteric conduit to transmit information on the RNA-occupancy of the C3H domain to the RING domain that transfers ubiquitin to its substrate. Interestingly, the short alpha-helix of the C3H(2) domain predicted to interact with the CH domain in MKRN3 has a dual role in this family of ZNFs, where it can interact with both RNA and proteins [71]. Thus, RNA and the CH domain in MKRNs could be competing for the same mutually exclusive binding site on the C3H(2) domain.

The zinc ligation site of the previously uncharacterized CH-domain offers clues about its function. It was previously suggested that the CH-domain with three cysteines and three histidines could be a ZNFs with a binuclear zinc cluster [33]. A Zn_2_Cys_6_ configuration occurs in the dimeric α-helical GAL4 transcription factor so it was inferred that the CH-domain might also bind DNA through its six potential ligands [33]. The annotation of the CH-domain as a DNA-binding module has persisted in several reviews on the makorin family proteins. Yet, our present works shows the CH-domain is a monomer with a ββα-fold that binds a single Zn^2+^ ion through a CCHC chelation site comprised of C266, C269, H285, and C289. This makes it unlikely that the CH-domain binds DNA, since this typically requires 2 to 3 or more ZNFs with ββα-folds and CCHH metal binding sites rather than a CCHC metal binding site [35]. The CCHC ligand set established in this work also rules out an RNA-binding function which is typically associated with CCCH [39], CCHC-knuckle [41], or RANBP2-type CCCC [36] ZNFs. The closest sequence motif to the CH-domain is the CCHC-finger [42] found in protein-binding ZNFs such as those from FOG [72], NEMO [73], or ZNF750 [42]. Yet, the CH-domain is a sub-optimal match for this family since it has a 15- rather than 12-residue spacing between the second and third zinc-coordinating residues and it is missing the conserved hydrophobic and aromatic residues of this motif. We therefore used the MOTIF2 server (https://www.genome.jp/tools/motif/MOTIF2.html accessed on 8 November 2025) to look for the query pattern C-x(2)-C-x(15)-H-x(3)-C with the search restricted to human sequences. We found some 29 hits including the four makorins, parts of RING domains in other E3-ligases that share this spacing, accidental hits to cysteine rich proteins, and some 3-5 hits of unknown structure where AF3 predicts a ZNF, suggesting this motif is more common than currently realized. Amongst the hits there was only one other example with a known structure (PDB codes 2MUR, 2MUQ): a UBZ2-type ubiquitin-binding ZNF corresponding to residues 140-180 of FAAP20 (Fanconi anemia core complex-associated protein 20) [64].

Given that the closest motif to the CH-domain binds ubiquitin, we wondered if ubiquitin could also be an interaction partner for MKRN3. We did an AF3 simulation for full-length MKRN3 with 6 Zn^2+^ ions to satisfy all the zinc-binding sites in the five ZNF domains, and also included a ubiquitin molecule as a potential binding partner in the simulation. The AF3 program predicts that ubiquitin binds to the center (I290-F301) of the long α-helix following the β-hairpin of the CH domain (Figs. 3F and S6). There are currently four types of ubiquitin-binding zinc (UBZ) fingers recognized [64]. Although each type has unique structural properties and they coordinate zinc with different CCHH or CCHC residue combinations, they can share common structural motifs. A common but not universal [74,75] theme for UBZs, is that ubiquitin is bound through a long α-helix at the C-terminus of the ZNF, following either a CCHC β-hairpin, as in the case of the UBZ2 from FAAP20 [64], or a CCCC β-knuckle like in RABEX-5 [74,76]. The α−helix from the ZNF usually binds to a distinct part of ubiquitin, in the cleft between β-strands 3 and 5 on the opposite side of the structure from the ubiquitin α-helix. The long α-helix from the ZNF is oriented parallel to strand β5 of ubiquitin [64,74]. All these features occur in the AF3-predicted complex of MKRN3 and ubiquitin (Figs. 3F and S6). Part of the AF3-predicted binding site for ubiquitin overlaps the C-terminus of the α-helix in our CH-domain structure, the remainder is in the α-helix extension past the domain boundary in our fragment.

We did similar AF3 simulations for the remainder of the MKRN family. The domain spacing of the C3H(2)-CH-RING tandem is conserved in all four makorin proteins [1,3] and AF3 predicts conserved interdomain interaction between the CH and preceding C3H domains. All four makorin proteins are predicted by AF3 to bind ubiquitin in a similar location on the α-helix of the CH-domain. As such, the putative interactions are suggested to be general across the makorin family and not just a feature of MKRN3. To test the reliability of the AF3 prediction that MK3 binds ubiquitin, we repeated the simulations with 9 random similarly sized proteins that have no sequence, structural, or functional relationship to ubiquitin (Fig. S7). All 9 decoy proteins bound to the C3H(2)-CH-RING segment of MKRN3, perhaps because this is the longest segment of predicted contiguous structure in the protein that is uninterrupted by disordered regions. About half of the decoys bound to the long helix segment at the end of the CH-domain, particularly the proteins with substantial β-sheet structure. All of the decoy proteins had similarly high AF3 pLDDT > 70 confidence scores, comparable to ubiquitin (Fig. S7). Thus, we used AF3 to test for ubiquitin binding to MKRN3 based on our hypothesis that the CH-domain is similar to a ubiquitin-binding ZNF. We could not verify or refute our hypothesis because AF3 generates comparably ‘reasonable’ structures for putative complexes between MKRN3 and randomly chosen proteins.

Our results highlight that AI structural predictions can be equivocal. Ultimately, experimental structural biology is still needed to establish if unusual ZNFs like MKRN3-CH adopt folded structures, to accurately determine nuances of the structure like the true length of the α-helix in the CH domain, and to verify the interactome of possible intra and inter-molecular association reactions.

## CRediT authorship contribution statement

A. **J. Rua:** Investigation, formal analysis, writing – original draft, writing-review and editing.

**A.T. Alexandrescu:** Conceptualization, resources, investigation, formal analysis, writing-review and editing, project administration.

## CONFLICT OF INTEREST

The authors declare no potential conflict of interest.

## FUNDING

NMR experiments were done at the Gregory P. Mullen NMR Structural Biology Facility of UConn Health, which is a member of the NSF Network for Advanced NMR (grants 1946970 and 2529058). 800 MHz and 700 MHz instruments were funded by NIH grants S10RR023041 and S10OD034297, respectively.

## ACKNOWLEDGEMENTS

We thank Prof. Carolyn Teschke for use of her CD and UV-Vis instruments

## SUPPLEMENTARY DATA

Supplementary data to this article can be found online at https://**????**

## Abbreviations

AF3, AlphaFold 3; CD, circular dichroism; C3H, zinc fingers with a ligand set consisting of three cysteines followed by a histidine that typically have RNA-binding functions; C3H(2)-CH-RING, three-domain segment of MKRN3 running between residues 238-365; CH-domain, cysteine/histidine-rich domain of makorins; CPP, central precocious puberty; GnRH, gonadotropin-releasing hormone; HPG, hypothalamic-pituitary-gonadal axis; MKRN, makorin; MKRN3, makorin-3; NMR, nuclear magnetic resonance; RING, (Really Interesting New Gene) zinc finger that is typically the active site of E3 ubiquitin-ligases featuring a binuclear zinc cluster ligated by C3HC4 ligands; UBZ, ubiquitin-binding zinc finger; ZNF, zinc finger

## Notes

https://www.rcsb.org/structure/unreleased/9P2Q

https://bmrb.io/data_library/held.shtml#53190

### Competing Interest Statement

The authors have declared no competing interest.

## REFERENCES

[1] L. Naule, U.B. Kaiser, Evolutionary Conservation of MKRN3 and Other Makorins and Their Roles in Puberty Initiation and Endocrine Functions, Semin Reprod Med. 37 (2019) 166–173. 10.1055/s-0039-3400965.

[2] T. Wang, W. Liu, C. Wang, X. Ma, M.F. Akhtar, Y. Li, L. Li, MRKNs: Gene, Functions, and Role in Disease and Infection, Front Oncol. 12 (2022) 862206. 10.3389/fonc.2022.862206.

[3] E.A. Guseva, M.A. Emelianova, V.N. Sidorova, A.N. Tyulpakov, O.A. Dontsova, P.V. Sergiev, Diversity of Molecular Functions of RNA-Binding Ubiquitin Ligases from the MKRN Protein Family, Biochemistry (Mosc). 89 (2024) 1558–1572. 10.1134/S0006297924090037.

[4] A. Bohne, A. Darras, H. D’Cotta, J.F. Baroiller, D. Galiana-Arnoux, J.N. Volff, The vertebrate makorin ubiquitin ligase gene family has been shaped by large-scale duplication and retroposition from an ancestral gonad-specific, maternal-effect gene, BMC Genomics. 11 (2010) 721. 10.1186/1471-2164-11-721.

[5] M. Jong, T.A. Gray, Y. Ji, C.C. Glenn, S. Saitoh, D.J. Driscoll, R.D. Nicholls, A novel imprinted gene, encoding a RING zinc-finger protein, and overlapping antisense transcript in the Prader–Willi syndrome critical region, Human Molecular Genetics. 8 (1999) 783–793. 10.1093/hmg/8.5.783.

[6] S. Palumbo, G. Cirillo, F. Aiello, A. Papparella, E. Miraglia Del Giudice, A. Grandone, MKRN3 role in regulating pubertal onset: the state of art of functional studies, Front Endocrinol (Lausanne). 13 (2022) 991322. 10.3389/fendo.2022.991322.

[7] M.F. Faienza, F. Urbano, L.A. Moscogiuri, M. Chiarito, S. De Santis, P. Giordano, Genetic, epigenetic and enviromental influencing factors on the regulation of precocious and delayed puberty, Front Endocrinol (Lausanne). 13 (2022) 1019468. 10.3389/fendo.2022.1019468.

[8] S. Vadakkadath Meethal, C.S. Atwood, The role of hypothalamic-pituitary-gonadal hormones in the normal structure and functioning of the brain, Cell Mol Life Sci. 62 (2005) 257–270. 10.1007/s00018-004-4381-3.

[9] Q. Xie, Y. Kang, C. Zhang, Y. Xie, C. Wang, J. Liu, C. Yu, H. Zhao, D. Huang, The Role of Kisspeptin in the Control of the Hypothalamic-Pituitary-Gonadal Axis and Reproduction, Front Endocrinol (Lausanne). 13 (2022) 925206. 10.3389/fendo.2022.925206.

[10] J. Liu, T. Li, M. Peng, M. Luo, Z. Gui, S. Long, Z. Mo, W. He, The Key Roles of Makorin RING Finger Protein 3 (MKRN3) During the Development of Pubertal Initiation and Central Precocious Puberty (CPP), Curr Mol Med. 23 (2023) 668–677. 10.2174/1566524022666220624105430.

[11] P. Abreu Ana, A. Dauber, B. Macedo Delanie, D. Noel Sekoni, N. Brito Vinicius, C. Gill John, P. Cukier, R. Thompson Iain, M. Navarro Victor, C. Gagliardi Priscila, T. Rodrigues, C. Kochi, A. Longui Carlos, D. Beckers, F. de Zegher, R. Montenegro Luciana, B. Mendonca Berenice, S. Carroll Rona, N. Hirschhorn Joel, C. Latronico Ana, A. Kaiser Ursula, Central Precocious Puberty Caused by Mutations in the Imprinted Gene MKRN3, New England Journal of Medicine. 368 (2013) 2467–2475. 10.1056/NEJMoa1302160.

[12] A.P. Abreu, C.A. Toro, Y.B. Song, V.M. Navarro, M.A. Bosch, A. Eren, J.N. Liang, R.S. Carroll, A.C. Latronico, O.K. Rønnekleiv, C.F. Aylwin, A. Lomniczi, S. Ojeda, U.B. Kaiser, MKRN3 inhibits the reproductive axis through actions in kisspeptin-expressing neurons, The Journal of Clinical Investigation. 130 (2020) 4486–4500. 10.1172/JCI136564.

[13] C. Li, W. Lu, L. Yang, Z. Li, X. Zhou, R. Guo, J. Wang, Z. Wu, Z. Dong, G. Ning, Y. Shi, Y. Gu, P. Chen, Z. Hao, T. Han, M. Yang, W. Wang, X. Huang, Y. Li, S. Gao, R. Hu, MKRN3 regulates the epigenetic switch of mammalian puberty via ubiquitination of MBD3, National Science Review. 7 (2020) 671–685. 10.1093/nsr/nwaa023.

[14] A.P. Abreu, D.B. Macedo, V.N. Brito, U.B. Kaiser, A.C. Latronico, A new pathway in the control of the initiation of puberty: the MKRN3 gene, J Mol Endocrinol. 54 (2015) R131-

15. 139. 10.1530/JME-14-0315.

[15] L. Maione, L. Naule, U.B. Kaiser, Makorin RING finger protein 3 and central precocious puberty, Curr Opin Endocr Metab Res. 14 (2020) 152–159. 10.1016/j.coemr.2020.08.003.

[16] J.C. Magnotto, A. Mancini, K. Bird, L. Montenegro, F. Tütüncüler, S.A. Pereira, V. Simas, L. Garcia, S.A. Roberts, D. Macedo, M. Magnuson, P. Gagliardi, N. Mauras, S.F. Witchel, R.S. Carroll, A.C. Latronico, U.B. Kaiser, A.P. Abreu, Novel MKRN3 Missense Mutations Associated With Central Precocious Puberty Reveal Distinct Effects on Ubiquitination, The Journal of Clinical Endocrinology & Metabolism. 108 (2023) 1646–1656. 10.1210/clinem/dgad151.

[17] A. Alghamdi, Precocious Puberty: Types, Pathogenesis and Updated Management, Cureus. 15 (2023) e47485. 10.7759/cureus.47485.

[18] M. Chen, E.A. Eugster, Central Precocious Puberty: Update on Diagnosis and Treatment, Paediatr Drugs. 17 (2015) 273–281. 10.1007/s40272-015-0130-8.

[19] A.C. Latronico, V.N. Brito, J.-C. Carel, Causes, diagnosis, and treatment of central precocious puberty, The Lancet Diabetes & Endocrinology. 4 (2016) 265–274. 10.1016/S2213-8587(15)00380-0.

[20] E.L. Zevin, E.A. Eugster, Central precocious puberty: a review of diagnosis, treatment, and outcomes, The Lancet Child & Adolescent Health. 7 (2023) 886–896. 10.1016/S2352-4642(23)00237-7.

[21] S.J. Lee, E.M. Yang, J.Y. Seo, C.J. Kim, Effects of gonadotropin-releasing hormone agonist therapy on body mass index and height in girls with central precocious puberty, Chonnam Med J. 48 (2012) 27–31. 10.4068/cmj.2012.48.1.27.

[22] P. Fanis, M. Morrou, M. Tomazou, K. Michailidou, G.M. Spyrou, M. Toumba, N. Skordis, V. Neocleous, L.A. Phylactou, Methylation status of hypothalamic Mkrn3 promoter across puberty, Front Endocrinol (Lausanne). 13 (2022) 1075341. 10.3389/fendo.2022.1075341.

[23] A.P. Abreu, C.A. Toro, Y.B. Song, V.M. Navarro, M.A. Bosch, A. Eren, J.N. Liang, R.S. Carroll, A.C. Latronico, O.K. Ronnekleiv, C.F. Aylwin, A. Lomniczi, S. Ojeda, U.B. Kaiser, MKRN3 inhibits the reproductive axis through actions in kisspeptin-expressing neurons, J Clin Invest. 130 (2020) 4486–4500. 10.1172/JCI136564.

[24] A. Dold, H. Han, N. Liu, A. Hildebrandt, M. Brüggemann, C. Rücklé, H. Hänel, A. Busch, P. Beli, K. Zarnack, J. König, J.-Y. Roignant, P. Lasko, Makorin 1 controls embryonic patterning by alleviating Bruno1-mediated repression of oskar translation, PLOS Genetics. 16 (2020) e1008581. 10.1371/journal.pgen.1008581.

[25] Z. Yu, X. Li, J. Huang, J. Pan, J. Cheng, P. Liu, M. Yang, T. Chen, Q. Zhang, Y. Zhou, J. Wu, T. Han, J. Li, Y. Xu, M. Wen, X. Zhang, C. Wang, X. Cao, The RNA-binding E3 ligase MKRN2 selectively disrupts Il6 translation to restrain inflammation, Nat Immunol. 26 (2025) 1036–1047. 10.1038/s41590-025-02183-x.

[26] C. Li, W. Lu, L. Yang, Z. Li, X. Zhou, R. Guo, J. Wang, Z. Wu, Z. Dong, G. Ning, Y. Shi, Y. Gu, P. Chen, Z. Hao, T. Han, M. Yang, W. Wang, X. Huang, Y. Li, S. Gao, R. Hu, MKRN3 regulates the epigenetic switch of mammalian puberty via ubiquitination of MBD3, Natl Sci Rev. 7 (2020) 671–685. 10.1093/nsr/nwaa023.

[27] C. Li, T. Han, Q. Li, M. Zhang, R. Guo, Y. Yang, W. Lu, Z. Li, C. Peng, P. Wu, X. Tian, Q. Wang, Y. Wang, V. Zhou, Z. Han, H. Li, F. Wang, R. Hu, MKRN3-mediated ubiquitination of Poly(A)-binding proteins modulates the stability and translation of GNRH1 mRNA in mammalian puberty, Nucleic Acids Res. 49 (2021) 3796–3813. 10.1093/nar/gkab155.

[28] A. Hildebrandt, M. Bruggemann, C. Ruckle, S. Boerner, J.B. Heidelberger, A. Busch, H. Hanel, A. Voigt, M.M. Mockel, S. Ebersberger, A. Scholz, A. Dold, T. Schmid, I. Ebersberger, J.Y. Roignant, K. Zarnack, J. Konig, P. Beli, The RNA-binding ubiquitin ligase MKRN1 functions in ribosome-associated quality control of poly(A) translation, Genome Biol. 20 (2019) 216. 10.1186/s13059-019-1814-0.

[29] J.C. Magnotto, A. Mancini, K. Bird, L. Montenegro, F. Tutunculer, S.A. Pereira, V. Simas, L. Garcia, S.A. Roberts, D. Macedo, M. Magnuson, P. Gagliardi, N. Mauras, S.F. Witchel, R.S. Carroll, A.C. Latronico, U.B. Kaiser, A.P. Abreu, Novel MKRN3 Missense Mutations Associated With Central Precocious Puberty Reveal Distinct Effects on Ubiquitination, J Clin Endocrinol Metab. 108 (2023) 1646–1656. 10.1210/clinem/dgad151.

[30] A. Hildebrandt, M. Brüggemann, C. Rücklé, S. Boerner, J.B. Heidelberger, A. Busch, H. Hänel, A. Voigt, M.M. Möckel, S. Ebersberger, A. Scholz, A. Dold, T. Schmid, I. Ebersberger, J.-Y. Roignant, K. Zarnack, J. König, P. Beli, The RNA-binding ubiquitin ligase MKRN1 functions in ribosome-associated quality control of poly(A) translation, Genome Biology. 20 (2019) 216. 10.1186/s13059-019-1814-0.

[31] R.L. Carpenedo, C.P. A., W.L. and Stanford, MKRN1: Uncovering function by an unbiased systems approach, Cell Cycle. 15 (2016) 303–304. 10.1080/15384101.2015.1124698.

[32] C. Garcia-Barcena, N. Osinalde, J. Ramirez, U. Mayor, How to Inactivate Human Ubiquitin E3 Ligases by Mutation, Front Cell Dev Biol. 8 (2020) 39. 10.3389/fcell.2020.00039.

[33] T.A. Gray, L. Hernandez, A.H. Carey, M.A. Schaldach, M.J. Smithwick, K. Rus, J.A. Marshall Graves, C.L. Stewart, R.D. Nicholls, The ancient source of a distinct gene family encoding proteins featuring RING and C(3)H zinc-finger motifs with abundant expression in developing brain and nervous system, Genomics. 66 (2000) 76–86. 10.1006/geno.2000.6199.

[34] The_UniProt_Consortium, UniProt: the Universal Protein Knoledgebase in 2023, Nucleic Acids Res. 51 (2023) D523–D531. 10.1093/nar/gku989.

[35] A.V. Persikov, M. Singh, De novo prediction of DNA-binding specificities for Cys2His2 zinc finger proteins, Nucleic Acids Res. 42 (2014) 97–108. 10.1093/nar/gkt890.

[36] R.E. Harris, A.J. Rua, A.T. Alexandrescu, Zinc-Induced Folding and Solution Structure of the Eponymous Novel Zinc Finger from the ZC4H2 Protein, Biomolecules. 15 (2025). 10.3390/biom15081091.

[37] J. Abramson, J. Adler, J. Dunger, R. Evans, T. Green, A. Pritzel, O. Ronneberger, L. Willmore, A.J. Ballard, J. Bambrick, S.W. Bodenstein, D.A. Evans, C.-C. Hung, M. O’Neill, D. Reiman, K. Tunyasuvunakool, Z. Wu, A. Žemgulytė, E. Arvaniti, C. Beattie, O. Bertolli, A. Bridgland, A. Cherepanov, M. Congreve, A.I. Cowen-Rivers, A. Cowie, M. Figurnov, F.B. Fuchs, H. Gladman, R. Jain, Y.A. Khan, C.M.R. Low, K. Perlin, A. Potapenko, P. Savy, S. Singh, A. Stecula, A. Thillaisundaram, C. Tong, S. Yakneen, E.D. Zhong, M. Zielinski, A. Žídek, V. Bapst, P. Kohli, M. Jaderberg, D. Hassabis, J.M. Jumper, Accurate structure prediction of biomolecular interactions with AlphaFold 3, Nature. 630 (2024) 493–500. 10.1038/s41586-024-07487-w.

[38] T. Chen, L. Chen, H. Wu, R. Xie, F. Wang, X. Chen, H. Sun, F. Xiao, Low Frequency of MKRN3 and DLK1 Variants in Chinese Children with Central Precocious Puberty, Int J Endocrinol. 2019 (2019) 9879367. 10.1155/2019/9879367.

[39] J.L. Michalek, A.N. Besold, S.L. Michel, Cysteine and histidine shuffling: mixing and matching cysteine and histidine residues in zinc finger proteins to afford different folds and function, Dalton Trans. 40 (2011) 12619–12632. 10.1039/c1dt11071c.

[40] W.M. Matousek, A.T. Alexandrescu, NMR structure of the C-terminal domain of SecA in the free state, Biochim Biophys Acta. 1702 (2004) 163–171. 10.1016/j.bbapap.2004.08.012.

[41] U. Aceituno-Valenzuela, R. Micol-Ponce, M.R. Ponce, Genome-wide analysis of CCHC-type zinc finger (ZCCHC) proteins in yeast, Arabidopsis, and humans, Cell Mol Life Sci. 77 (2020) 3991–4014. 10.1007/s00018-020-03518-7.

[42] A.J. Rua, R.D. Whitehead, A.T. Alexandrescu, NMR structure verifies the eponymous zinc finger domain of transcription factor ZNF750, Journal of Structural Biology: X. 8 (2023) 100093. 10.1016/j.yjsbx.2023.100093.

[43] R.E. Harris, R.D. Whitehead Iii, A.T. Alexandrescu, Solution structure of the Z0 domain from transcription repressor BCL11A sheds light on the sequence properties of protein-binding zinc fingers, Protein Science. 34 (2025) e70097. 10.1002/pro.70097.

[44] J.M. Walker, The Bicinchoninic Acid (BCA) Assay for Protein Quantitation, in: J.M. Walker (Ed.), The Protein Protocols Handbook, Humana Press, Totowa, NJ, 2009, pp. 11-15.

[45] E. Ivanova, M. Ball, H. Lu, Zinc binding of Tim10: Evidence for existence of an unstructured binding intermediate for a zinc finger protein, 71 (2008) 467–475. 10.1002/prot.21713.

[46] A.J. Rua, A.T. Alexandrescu, Formerly degenerate seventh zinc finger domain from transcription factor ZNF711 rehabilitated by experimental NMR structure, Protein Science. 33 (2024) e5149. 10.1002/pro.5149.

[47] K. Kluska, J. Adamczyk, A. Krezel, Metal binding properties, stability and reactivity of zinc fingers, Coordination Chemistry Reviews. 367 (2018) 18–64.

[48] T.R. Young, Z. Xiao, Principles and practice of determining metal-protein affinities, Biochem J. 478 (2021) 1085–1116. 10.1042/BCJ20200838.

[49] R.D. Whitehead, 3rd, C.M. Teschke, A.T. Alexandrescu, Pulse-field gradient nuclear magnetic resonance of protein translational diffusion from native to non-native states, Protein Sci. 31 (2022) e4321. 10.1002/pro.4321.

[50] D.S. Wishart, C.G. Bigam, J. Yao, F. Abildgaard, H.J. Dyson, E. Oldfield, J.L. Markley, B.D. Sykes, 1H, 13C and 15N chemical shift referencing in biomolecular NMR, Journal of Biomolecular NMR. 6 (1995) 135-140. 10.1007/BF00211777.

[51] C. Harprecht, O. Okifo, K.J. Robbins, T. Motwani, A.T. Alexandrescu, C.M. Teschke, Contextual Role of a Salt Bridge in the Phage P22 Coat Protein I-Domain*, Journal of Biological Chemistry. 291 (2016) 11359–11372. 10.1074/jbc.M116.716910.

[52] K. Wuthrich, M. Billeter, W. Braun, Pseudo-structures for the 20 common amino acids for use in studies of protein conformations by measurements of intramolecular proton-proton distance constraints with nuclear magnetic resonance, J Mol Biol. 169 (1983) 949–961. 10.1016/s0022-2836(83)80144-2.

[53] Y. Shen, A. Bax, Protein structural information derived from NMR chemical shift with the neural network program TALOS-N, Methods Mol Biol. 1260 (2015) 17–32. 10.1007/978-1-4939-2239-0_2.

[54] S. Ramboarina, S. Druillennec, N. Morellet, S. Bouaziz, B.P. Roques, Target Specificity of Human Immunodeficiency Virus Type 1 NCp7 Requires an Intact Conformation of Its CCHC N-Terminal Zinc Finger, Journal of Virology. 78 (2004) 6682–6687. 10.1128/jvi.78.12.6682-6687.2004.

[55] G.A. Bermejo, N. Tjandra, G.M. Clore, C.D. Schwieters, Xplor-NIH: Better parameters and protocols for NMR protein structure determination, Protein Science. 33 (2024) e4922. 10.1002/pro.4922.

[56] W. Rieping, M. Habeck, B. Bardiaux, A. Bernard, T.E. Malliavin, M. Nilges, ARIA2: Automated NOE assignment and data integration in NMR structure calculation, Bioinformatics. 23 (2007) 381–382. 10.1093/bioinformatics/btl589.

[57] M.W. Maciejewski, A.D. Schuyler, M.R. Gryk, I.I. Moraru, P.R. Romero, E.L. Ulrich, H.R. Eghbalnia, M. Livny, F. Delaglio, J.C. Hoch, NMRbox: A Resource for Biomolecular NMR Computation, Biophysical Journal. 112 (2017) 1529–1534. 10.1016/j.bpj.2017.03.011.

[58] Y. Wang, Y. Yu, Y. Pang, H. Yu, W. Zhang, X. Zhao, J. Yu, The distinct roles of zinc finger CCHC-type (ZCCHC) superfamily proteins in the regulation of RNA metabolism, RNA Biol. 18 (2021) 2107–2126. 10.1080/15476286.2021.1909320.

[59] J.M. Berg, Y. Shi, The galvanization of biology: a growing appreciation for the roles of zinc, Science. 271 (1996) 1081–1085. 10.1126/science.271.5252.1081.

[60] A. Klug, The discovery of zinc fingers and their applications in gene regulation and genome manipulation, Annu Rev Biochem. 79 (2010) 213–231. 10.1146/annurev-biochem-010909-095056.

[61] A. Nomura, Y. Sugiura, Contribution of individual zinc ligands to metal binding and peptide folding of zinc finger peptides, Inorg Chem. 41 (2002) 3693–3698. 10.1021/ic025557p.

[62] V. Sivo, G. D’Abrosca, L. Russo, R. Iacovino, P.V. Pedone, R. Fattorusso, C. Isernia, G. Malgieri, Co(II) Coordination in Prokaryotic Zinc Finger Domains as Revealed by UV-Vis Spectroscopy, Bioinorg Chem Appl. 2017 (2017) 1527247. 10.1155/2017/1527247.

[63] A.J. Rua, R.D. Whitehead, 3rd, A.T. Alexandrescu, NMR structure verifies the eponymous zinc finger domain of transcription factor ZNF750, J Struct Biol X. 8 (2023) 100093. 10.1016/j.yjsbx.2023.100093.

[64] J.L. Wojtaszek, S. Wang, H. Kim, Q. Wu, A.D. D’Andrea, P. Zhou, Ubiquitin recognition by FAAP20 expands the complex interface beyond the canonical UBZ domain, Nucleic Acids Res. 42 (2014) 13997–14005. 10.1093/nar/gku1153.

[65] T.R. Alderson, I. Pritisanac, D. Kolaric, A.M. Moses, J.D. Forman-Kay, Systematic identification of conditionally folded intrinsically disordered regions by AlphaFold2, Proc Natl Acad Sci U S A. 120 (2023) e2304302120. 10.1073/pnas.2304302120.

[66] P.S. Kim, R.L. Baldwin, Specific intermediates in the folding reactions of small proteins and the mechanism of protein folding, Annu Rev Biochem. 51 (1982) 459–489. 10.1146/annurev.bi.51.070182.002331.

[67] D.A. Dolgikh, R.I. Gilmanshin, E.V. Brazhnikov, V.E. Bychkova, G.V. Semisotnov, S. Venyaminov, O.B. Ptitsyn, Alpha-Lactalbumin: compact state with fluctuating tertiary structure?, FEBS Lett. 136 (1981) 311–315. 10.1016/0014-5793(81)80642-4.

[68] L.J. Smith, A.T. Alexandrescu, M. Pitkeathly, C.M. Dobson, Solution structure of a peptide fragment of human alpha-lactalbumin in trifluoroethanol: a model for local structure in the molten globule, Structure. 2 (1994) 703–712. 10.1016/s0969-2126(00)00071-x.

[69] T. Miura, T. Satoh, H. Takeuchi, Role of metal-ligand coordination in the folding pathway of zinc finger peptides, Biochim Biophys Acta. 1384 (1998) 171–179. 10.1016/s0167-4838(98)00015-6.

[70] W. Li, J. Zhang, J. Wang, W. Wang, Metal-coupled folding of Cys2His2 zinc-finger, J Am Chem Soc. 130 (2008) 892–900. 10.1021/ja075302g.

[71] M. Teplova, D.J. Patel, Structural insights into RNA recognition by the alternative-splicing regulator muscleblind-like MBNL1, Nat Struct Mol Biol. 15 (2008) 1343–1351. 10.1038/nsmb.1519.

[72] J.M. Matthews, K. Kowalski, C.K. Liew, B.K. Sharpe, A.H. Fox, M. Crossley, J.P. MacKay, A class of zinc fingers involved in protein-protein interactions biophysical characterization of CCHC fingers from fog and U-shaped, Eur J Biochem. 267 (2000) 1030–1038. 10.1046/j.1432-1327.2000.01095.x.

[73] F. Cordier, E. Vinolo, M. Veron, M. Delepierre, F. Agou, Solution structure of NEMO zinc finger and impact of an anhidrotic ectodermal dysplasia with immunodeficiency-related point mutation, J Mol Biol. 377 (2008) 1419–1432. 10.1016/j.jmb.2008.01.048.

[74] S. Lee, Y.C. Tsai, R. Mattera, W.J. Smith, M.S. Kostelansky, A.M. Weissman, J.S. Bonifacino, J.H. Hurley, Structural basis for ubiquitin recognition and autoubiquitination by Rabex-5, Nat Struct Mol Biol. 13 (2006) 264–271. 10.1038/nsmb1064.

[75] A.A. Rizzo, P.E. Salerno, I. Bezsonova, D.M. Korzhnev, NMR structure of the human Rad18 zinc finger in complex with ubiquitin defines a class of UBZ domains in proteins linked to the DNA damage response, Biochemistry. 53 (2014) 5895–5906. 10.1021/bi500823h.

[76] L. Penengo, M. Mapelli, A.G. Murachelli, S. Confalonieri, L. Magri, A. Musacchio, P.P. Di Fiore, S. Polo, T.R. Schneider, Crystal structure of the ubiquitin binding domains of rabex-5 reveals two modes of interaction with ubiquitin, Cell. 124 (2006) 1183–1195. 10.1016/j.cell.2006.02.020.

