## Supporting Information for "Solution Structure of the Novel CH-domain zinc finger from the puberty regulator Makorin-3"

**Table of contents:**

**Figure S1 - $CoCl_2$ binding curve for the CH domain**
**Figure S2 - DOSY spectrum of $Zn^{2+}$-bound MKRN3-CH**
**Figure S3 - Wüthrich secondary structure diagram of $Zn^{2+}$-bound MKRN3-CH**
**Figure S4 - NMR temperature titration of $Zn^{2+}$-bound MKRN3-CH**
**Figure S5 - High-temperature CD spectra of $Zn^{2+}$-bound MKRN3-CH.**
**Figure S6 - AF3-predicted interactions between MKRN3 domains and ubiquitin**
**Figure S7 - AF3 predictions of MKRN3 complexes with various proteins**

**Table S1 - Statistics for the 20 best NMR structures of MKRN3-CH (PDB 9P2Q).**

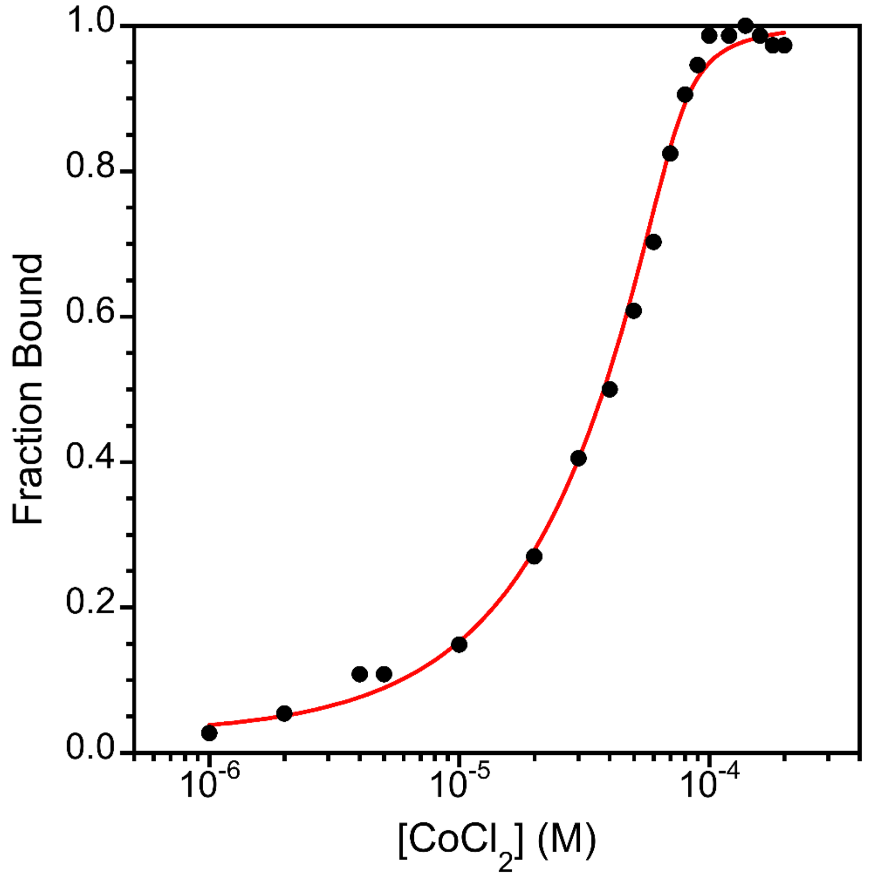

**Supplementary Figure S1 $CoCl_2$ binding curve for the CH domain.** The CH peptide concentration was 75 μM in 10 mM Tris pH 7.0 containing 0.5 mM of the reducing agent TCEP. The sample temperature was 25 °C. The added $CoCl_2$ concentration was varied from 1 to 200 μM and is shown on a semi-logarithmic X-axis. The fraction bound (Y-axis) was calculated from the d-d transition band at $A_{650}$ (Fig. 1F) Data were fitted to Eq. 1 of the main text to obtain a $K_d$ for $Co^{2+}$-binding of 1.8 x $10^{-6}$ M.

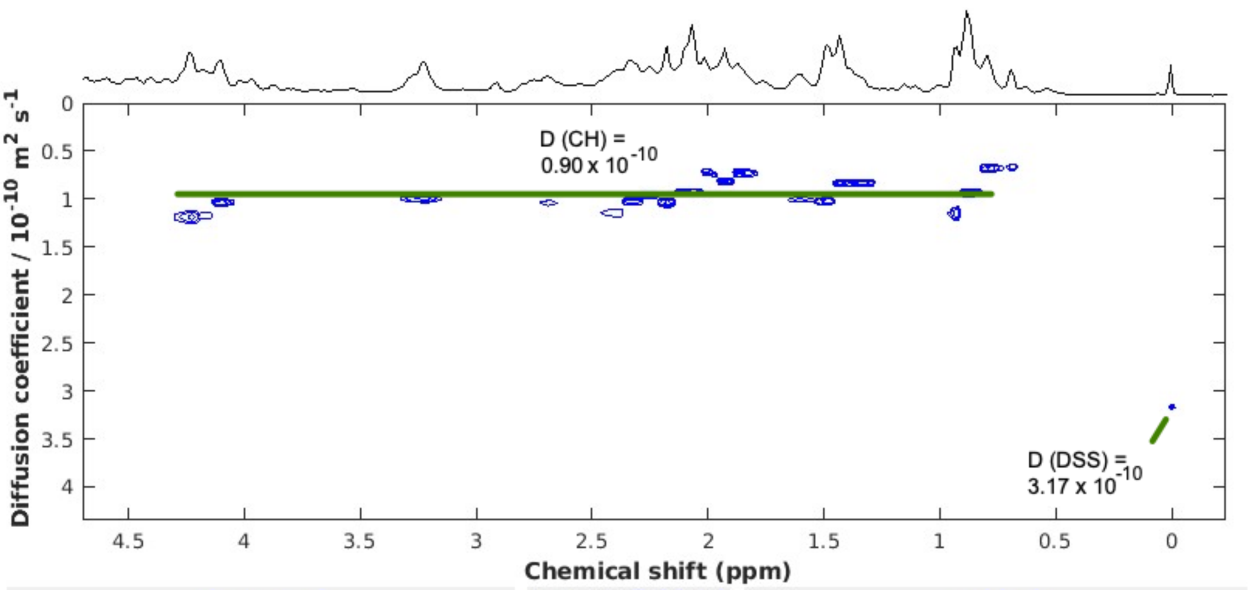

**Supplementary Figure S2 DOSY spectrum of $Zn^{2+}$-bound MKRN3-CH.** DOSY experiments were done with the Varian *Doneshot* pulse sequence [1] at a temperature of 25 °C and processed with the GNAT program [2] on the NMRbox platform [3]. The sample was 1.9 mM in MKRN3-CH, 2.3 mM in $ZnCl_2$ at pH 6.0 in 270 µL of 90% $H_2O$/10% $D_2O$ contained in a Shigemi microcell. The diffusion coefficients indicated in the plot were obtained for the DSS standard and the CH peptide. Using the $R_h$ (radius of hydration) value of 3.34 Å for DSS and the formula $R_{h,CH} = (D_{DSS}/D_{Z0})*R_{h,DSS}$, an Rh value of 11.8 Å was calculated for the folded $Zn^{2+}$-bound CH peptide, consistent with a monomer. From the empirical relationship for folded proteins $R_h = 4.75N^{0.29}$, where N is the number of amino acids [4], an $R_h$ of 12.86 Å is expected for a 31-residue monomer, and 15.72 Å for a 62-residue dimer.

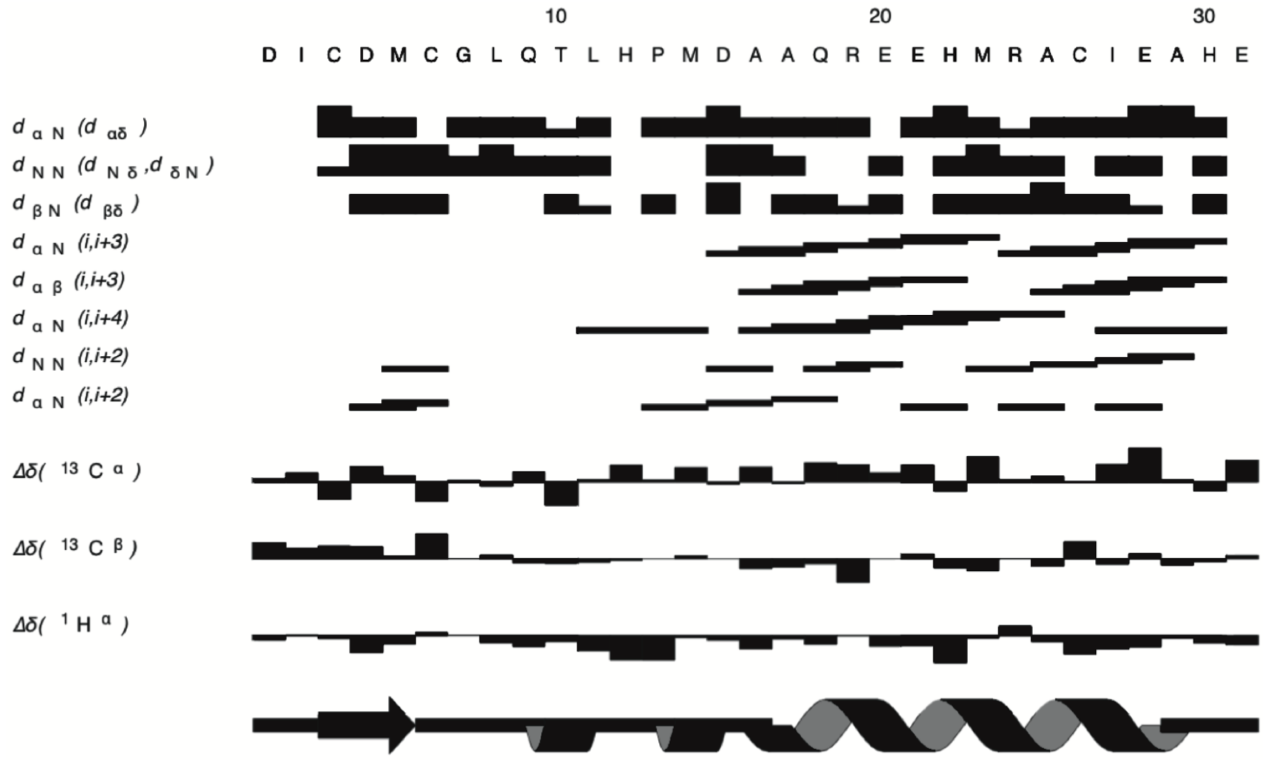

**Supplementary Figure S3 Wüthrich secondary structure diagram of $Zn^{2+}$-bound MKRN3-CH.** Short range NOEs (the sizes of the NOEs are indicated by the thickness of the bars) and $\Delta\delta$ chemical shift indices were used to calculate the consensus secondary structure shown at the bottom of the figure with the program DANGLE [5].

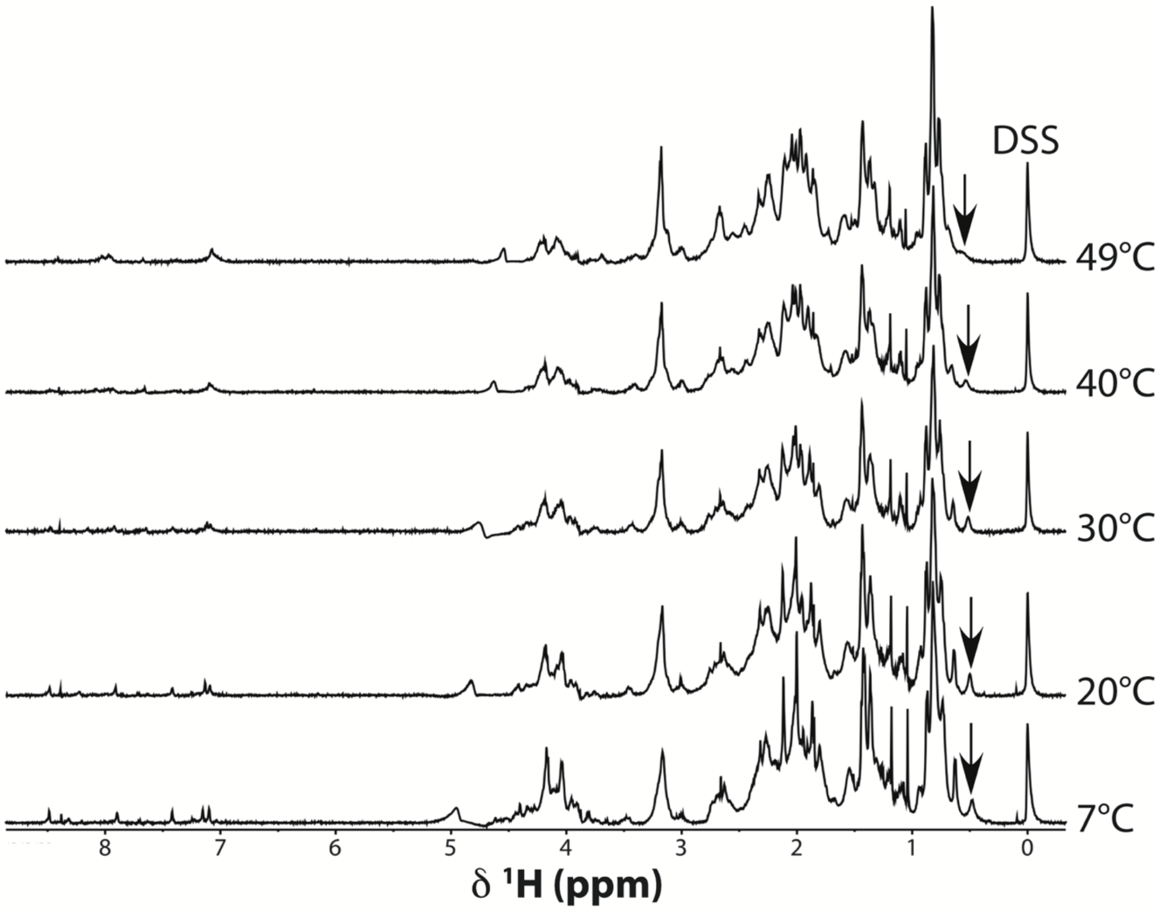

**Supplementary Figure S4 NMR temperature titration of $Zn^{2+}$-bound MKRN3-CH.** The full NMR spectra corresponding to the expansions in Fig. 4A are shown. The L11δa peak used for quantification is indicated by the black arrow.

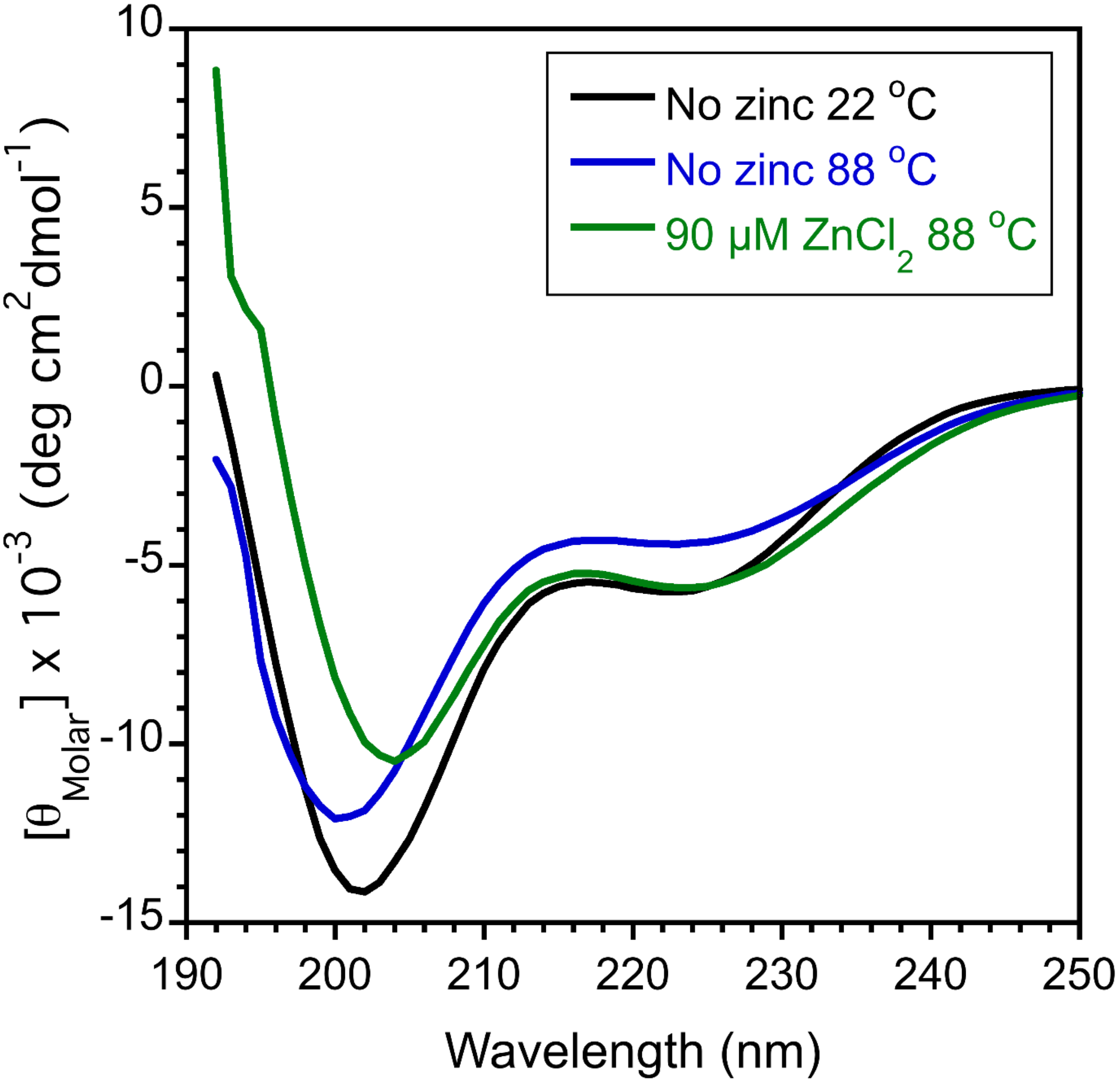

**Supplementary Figure S5 High-temperature CD spectra of $Zn^{2+}$-bound MKRN3-CH.** Black – spectrum of the apo peptide without zinc at 22 °C. Blue– spectrum of the apo peptide without zinc at 88 °C. Differences between the spectra of the apo peptide between low and high temperatures include an increase in ellipticity at 202 nm together with a small blue-shift from 202 to 200 nm at 88 °C, and a smaller increase in ellipticity at 222 nm at 88 °C. These changes could be temperature effects on the spectrum or a melting out of a small amount of residual structure between 22 and 88 °C. When $ZnCl_2$ is added to the peptide at 88 °C (green) there are small changes in the spectrum (red shift between 200 and 205 nm and a decrease in ellipticity at 222 nm) indicating the CH-domain still binds some $Zn^{2+}$ at 88 °C.

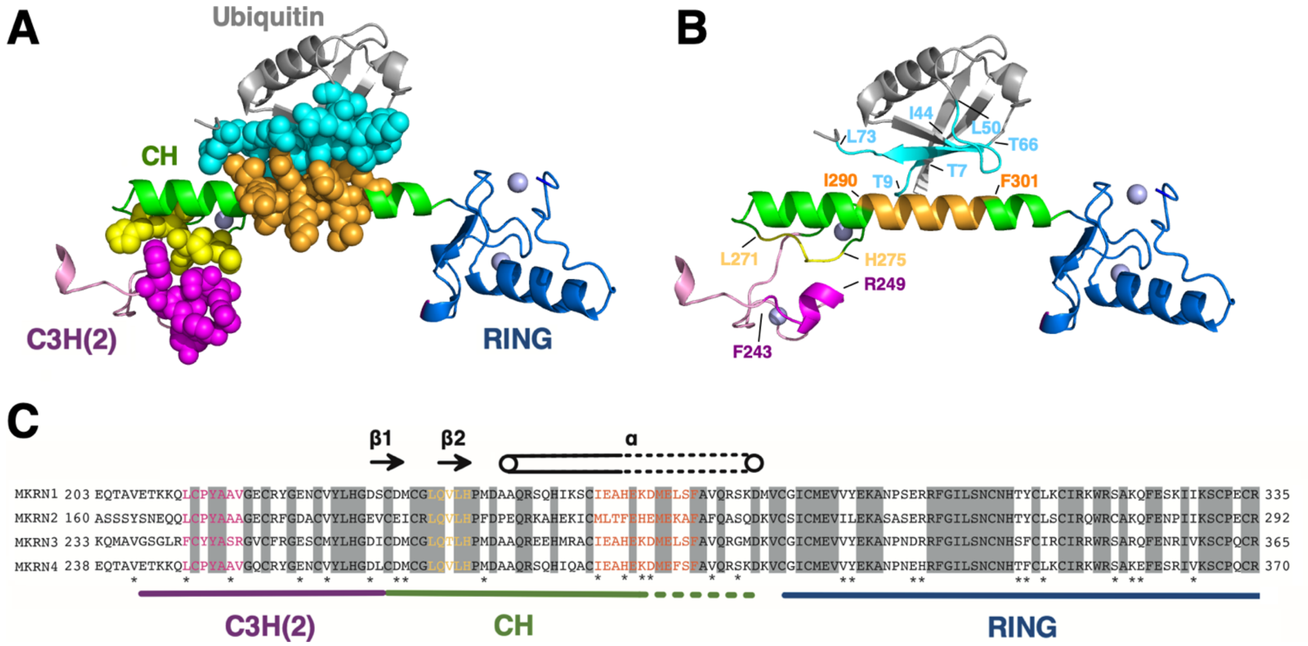

**Supplementary Figure S6 AF3-predicted interactions between MKRN3 domains and ubiquitin.** (**A**) Ribbon diagram of the C3H(2)-CH-RING segment from an AF3 prediction of the full-length MKRN3 protein bound to ubiquitin (see also Fig. 3F). To highlight the intramolecular interaction between the C3H(2) and CH domain and intermolecular interaction between the CH domain and human ubiquitin, the heavy atoms of the residues involved are shown as spheres. (**B**) Same as A but omitting the spheres to highlight the segments involved in the putative interactions: F243-R429 from the helix of the C3H(2) domain, L271-M277 from strand β2 of the CH domain, and I290-F301 from the extended helix of the CH domain. The segments interacting with MKRN3 from ubiquitin are T7-T9 (after β1), I44-L50 (after β3) and T66-L73 (β5). (**C**) Sequence alignment for the C3H(2)-CH-RING segment in the four makorins. The domain boundaries and secondary structure of the CH domain are shown. The dotted part of the α-helix is the extension predicted by AF3 that runs past the C-terminus of our peptide at E294 to about D308. Segments of the sequences corresponding to interacting residues (spheres in panel A) are indicated by 1-letter amino acid codes with the same color scheme as in panel A. Residues highlighted in grey are fully conserved across all four makorin family proteins. Residues with an asterisk below them have conserved properties but are not identical residues.

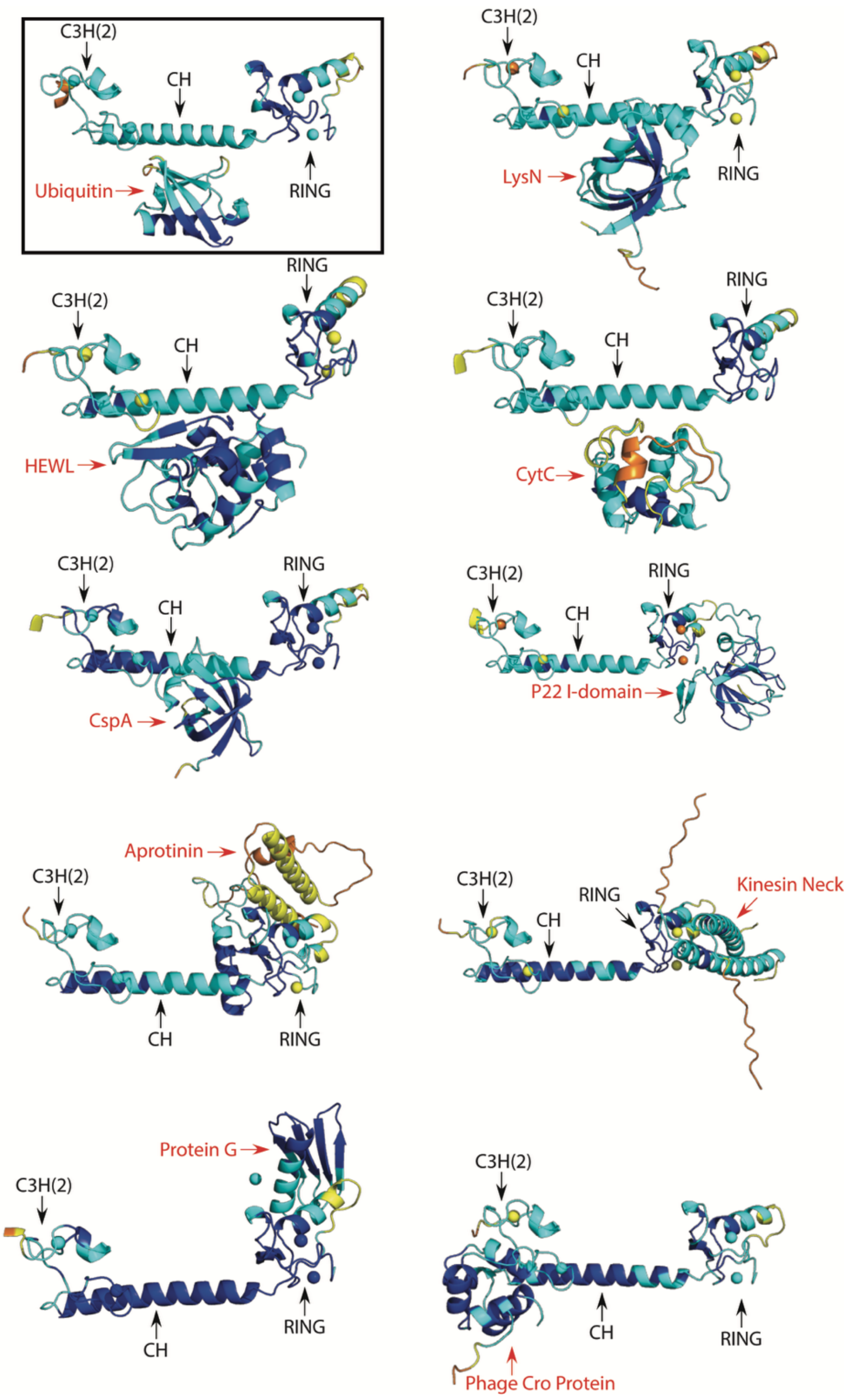

**Supplementary Figure S7  AF3 predictions of MKRN3 complexes with various proteins.** In each case the partner protein labeled in red was predicted to interact with the full-length 507 a.a. MKRN3 protein complexed with six $Zn^{2+}$ ions. Since each of the partner proteins bound to the C3H(2)-CH-RING segment (residues 238-365) only that region of the MKRN3 prediction is shown. The prediction for ubiquitin used in Figs. 3F and S6 is highlighted with a black rectangle in the upper left-hand corner. Colors are AF3 confidence score intervals: dark blue = pLDTT>90, cyan = 90>pLDDT>70, yellow = 70>pLDDT>50, orange = 50>pLDDT.

| **Table S1: Statistics for the 20 best NMR structures of MKRN3-CH (PDB 9P2Q)** | | |
|---|---|---|
| Restraints (total) | 372 | |
| Distance | 277 | |
| Intra-residue (\|i-j\|=0) | 97 | |
| Sequential (\|i-j\|=1) | 80 | |
| Medium Range (5>\|i-j\|>1) | 62 | |
| Long Range (\|i-j\|>5) | 38 | |
| Dihedral (25$\phi$, 25$\psi$, 8 $\chi$1) | 58 | |
| Hydrogen Bond | 20 | |
| Restraints to $Zn^{2+}$ atom | 8 | |
| $Zn^{2+}$ Coordination Geometry | 9 | |
| Restraint Violations | | |
| >0.5Å | 0 | |
| >5º | 0 | |
| Residual Restraint Violations | | |
| Distance (Å) | 0.035 +/- 0.004 | |
| Dihedral (º) | 0.66 +/- 0.17 | |
| RMS Deviations from ideal geometry | | |
| Bond lengths (Å) | 0.0039 +/- 0.0003 | |
| Bond angles (º) | 0.61 +/- 0.03 | |
| Improper Torsions (º) | 2.01 +/- 0.24 | |
| Ramachandran PROCHECK statistics for ordered residues | | |
| Residues in most favored regions | 87.7 % | |
| Residues in allowed regions | 11.7 % | |
| Residues in generously allowed regions | 0.6 % | |
| Residues in disallowed regions | 0.0 % | |
| Coordinate rms deviations (Å) | Backbone (Cα, N, C') | All heavy |
| NMR ensemble (1-31) to mean | 0.72 +/- 0.24 | 1.46 +/- 0.26 |
| NMR ensemble ordered (2-31) to mean | 0.65 +/- 0.26 | 1.38 +/- 0.24 |

[1] M.D. Pelta, G.A. Morris, M.J. Stchedroff, S.J. Hammond, A one-shot sequence for high-resolution diffusion-ordered spectroscopy, Magnetic Resonance in Chemistry. 40 (2002) S147-S152.

[2] L. Castanar, G.D. Poggetto, A.A. Colbourne, G.A. Morris, M. Nilsson, The GNAT: A new tool for processing NMR data, Magn Reson Chem. 56 (2018) 546-558. https://doi.org/10.1002/mrc.4717.

[3] M.W. Maciejewski, A.D. Schuyler, M.R. Gryk, I.I. Moraru, P.R. Romero, E.L. Ulrich, H.R. Eghbalnia, M. Livny, F. Delaglio, J.C. Hoch, NMRbox: A Resource for Biomolecular NMR Computation, Biophysical Journal. 112 (2017) 1529-1534. https://doi.org/https://doi.org/10.1016/j.bpj.2017.03.011.

[4] D.K. Wilkins, S.B. Grimshaw, V. Receveur, C.M. Dobson, J.A. Jones, L.J. Smith, Hydrodynamic radii of native and denatured proteins measured by pulse field gradient NMR techniques, Biochemistry. 38 (1999) 16424-16431. https://doi.org/10.1021/bi991765q.

[5] M.S. Cheung, M.L. Maguire, T.J. Stevens, R.W. Broadhurst, DANGLE: A Bayesian inferential method for predicting protein backbone dihedral angles and secondary structure, J Magn Reson. 202 (2010) 223-233. https://doi.org/10.1016/j.jmr.2009.11.008.
